# The vibriophage-encoded inhibitor OrbA abrogates BREX-mediated defense through the ATPase BrxC

**DOI:** 10.1101/2024.05.09.593382

**Authors:** Reid T Oshiro, Drew T Dunham, Kimberley D Seed

## Abstract

Bacteria and phages are locked in a co-evolutionary arms race where each entity evolves mechanisms to restrict the proliferation of the other. Phage-encoded defense inhibitors have proven powerful tools to interrogate how defense systems function. A relatively common defense system is BREX (Bacteriophage exclusion); however, how BREX functions to restrict phage infection remains poorly understood. A BREX system encoded by the SXT integrative and conjugative element, *Vch*Ind5, was recently identified in *Vibrio cholerae*, the causative agent of the diarrheal disease cholera. The lytic phage ICP1 that co-circulates with *V. cholerae* encodes the BREX inhibitor OrbA, but how OrbA inhibits BREX is unclear. Here, we determine that OrbA inhibits BREX using a unique mechanism from known BREX inhibitors by directly binding to the BREX component BrxC. BrxC has a functional ATPase domain that, when mutated, not only disrupts BrxC function but also alters how BrxC multimerizes. Furthermore, we find that OrbA binding disrupts BrxC-BrxC interactions. We determine that OrbA cannot bind BrxC encoded by the distantly related BREX system encoded by the SXT *Vch*Ban9, and thus fails to inhibit this BREX system that also circulates in epidemic *V. cholerae*. Lastly, we find that homologs of the *Vch*Ind5 BrxC are more diverse than the homologs of the *Vch*Ban9 BrxC. These data provide new insight into the function of the BrxC ATPase and highlight how phage-encoded inhibitors can disrupt phage defense systems using different mechanisms.

**Importance:** With renewed interest in phage therapy to combat antibiotic-resistant pathogens, understanding the mechanisms bacteria use to defend themselves against phages and the counter-strategies phages evolve to inhibit defenses is paramount. Bacteriophage exclusion (BREX) is a common defense system with few known inhibitors. Here, we probe how the vibriophage-encoded inhibitor OrbA inhibits the BREX system of *Vibrio cholerae*, the causative agent of the diarrheal disease cholera. By interrogating OrbA function, we have begun to understand the importance and function of a BREX component. Our results demonstrate the importance of identifying inhibitors against defense systems, as they are powerful tools for dissecting defense activity and can inform strategies to increase the efficacy of some phage therapies.

## Introduction

Bacteria are constantly antagonized by their viral counterparts, bacteriophages (herein phages). To restrict phage predation, bacteria use an arsenal of phage defense systems such as restriction-modification (R-M) systems and CRISPR-Cas^1–4^. In recent years, there has been a significant increase in the number of identified phage defense systems due to bioinformatic analysis and functional screens^5–9^. While the number of known defense systems has increased exponentially, the number of known phage-encoded inhibitors against these systems has lagged. A challenge with identifying phage-encoded inhibitors is that many are small proteins that lack sequence or structural similarity. Phage-encoded inhibitors are an important driver of the diversification of phage defense systems. For example, CRISPR-Cas systems, which are found in 40% of bacterial genomes, are extremely diverse, necessitating their classification into two classes subdivided into six types, which are further divided into more than 30 subtypes^4,10–12^. Both phages and other mobile elements can counter CRISPR-Cas mediated defense by encoding anti-CRISPR proteins (Acrs)^13,14^. There are currently over 100 Acrs, which boast a remarkable diversity of inhibitory mechanisms^13,14^. Arcs typically show high specificity towards particular variants of CRISPR-Cas, and thus, these inhibitors are a likely driving force for the diversification of CRISPR-Cas systems and the accumulation of other defenses in bacterial genomes. Additionally, inhibitors are powerful molecular tools that have been used to dissect the mechanisms of phage restriction by defense systems like CRISPR-Cas^13^. Acrs represent the largest family of known phage defense inhibitors; for all other known defense systems, few to no inhibitors have been identified.

The phage defense system BREX (bacteriophage exclusion) is a common defense system found in 8-10% of bacterial genomes^5,15^. Six types of BREX systems have been defined based on the phylogeny of the conserved alkaline phosphatase BrxZ (also known as PglZ)^15^, with the most common being type I comprised of six open reading frames, *brxABCXZL*. Some type I BREX systems encode additional genes, such as *brxR*, which encodes a transcriptional repressor, and *brxU*, which encodes a Type IV restriction enzyme^15–20^. Like R-M systems, BREX epigenetically modifies the host chromosome, and this modification is predicted to be used to distinguish self from foreign unmodified DNA^15,16^. Modification of the BREX system is coordinated by the DNA methyltransferase BrxX; however, BrxX alone is not sufficient for DNA modification, suggesting that BREX functions as a larger complex^16^. The structure of BrxA has been solved and has been shown to have DNA binding activity^21^, whereas BrxB has no known structure or domain features. BrxC has an ATPase domain that can be identified bioinformatically^15^. BrxL was previously predicted to have a Lon-like domain; however, a recent 3D structure of BrxL from *Acinetobacter* sp. 394 solved by cryo-electron microscopy shows BrxL has structural similarity to the MCM replicative helicase from archaea and eukaryotes^22^. How the BREX components function together to modulate modification and restriction is poorly understood. Type I BREX systems inhibit phage genome replication, however, degradation of the phage genome has not been observed^15,16^. How BREX restricts phage genome replication without degrading the phage genome is unclear. While BREX is a common phage defense, only three inhibitors are known to abrogate BREX activity: Ocr^23^, SAM lyase^24^, and OrbA^25^.

The inhibitor Ocr encoded by the *Escherichia coli* phage T7 was the first phage-encoded protein identified to abrogate BREX-mediated defense^23^. Ocr is also known to inhibit R-M systems by functioning as a DNA mimic ^26–28^. Similar to inhibiting R-M systems, Ocr directly binds the DNA methylase BrxX and restores T7 plaque formation^23^. The *E. coli* phage T3 inhibits BREX through the protein SAM lyase that degrades S-adenosyl Methionine (SAM) and inhibits the synthesis of SAM by binding to the synthase MetK, suggesting that SAM is important for BREX-mediated defense^24^. Like Ocr, SAM lyase has also been shown to provide protection against R-M systems^29^, suggesting there are common mechanisms used by R-M and BREX that permit these inhibitors to function broadly. While Ocr and SAM lyase inhibit BREX through different mechanisms, both have been powerful tools in dissecting how BREX functions and support the importance of BrxX in restricting phage. The third inhibitor, OrbA (<u>o</u>vercome restriction by <u>B</u>REX), functions through an unknown mechanism to protect the lytic phage ICP1 from BREX-mediated restriction in its host, the facultative pathogen *Vibrio cholerae*^25^.

OrbA was previously shown to protect ICP1 against the type I BREX system encoded by the integrative and <u>c</u>onjugative <u>e</u>lement (ICE), SXT *Vch*Ind5^25^. The SXT family of ICEs was initially discovered for harboring resistance to <u>s</u>ulfametho<u>x</u>azole and trimethoprim (SXT) and has recently been shown to encode various phage defense systems in a region known to vary in genomic content, denoted as hotspot 5^25,30^. OrbA does not share predicted structural similarity with known inhibitors, suggesting it inhibits BREX through a different mechanism^25^. While OrbA has been shown to inhibit the BREX system encoded by *Vch*Ind5 (BREX^Ind5^), OrbA does not inhibit the diverse type I BREX system encoded by the SXT *Vch*Ban9 (BREX^Ban9^), which is also encoded by epidemic *V. cholerae* strains^25^. The differences in OrbA sensitivity suggest that selective pressure to diversify the types of BREX systems in the epidemic *V. cholerae* population is important for protection from diverse phages encoding inhibitors against the BREX system.

Here, we interrogate how OrbA inhibits BREX^Ind5^. We demonstrate that BREX^Ind5^ inhibits ICP1 genome replication early in the replication cycle and does not appear to reduce the amount of phage genome present in the infected cell. We find that OrbA restores ICP1 genome replication to near wild-type levels by directly interacting with the BREX component, BrxC, supporting a new mechanism for inhibiting BREX-mediated defense. BrxC has an ATPase domain that is required for function as mutations that disrupt the ability to bind ATP or hydrolyze ATP cannot complement a *brxC* deletion. Additionally, proficiency in ATP binding and hydrolysis appears necessary for BrxC to interact with itself and most likely form a larger multimer. We find that the binding of OrbA alters how BrxC interacts with itself and suggests that OrbA disrupts BREX activity by abrogating BrxC multimerization. Furthermore, we determine that OrbA fails to inhibit BREX^Ban9^ because OrbA cannot interact with its respective BrxC. Homologs of BrxC proteins similar to the *Vch*Ban9 BrxC are highly similar, whereas homologs of the *Vch*Ind5 BrxC are highly diverse and appear to have been horizontally acquired. The diversification of BrxC proteins could explain how OrbA evolved to be a specific inhibitor to BREX^Ind5^, in comparison to the other known BREX inhibitors, which can also inhibit R-M systems. These findings help with deciphering the function of the proteins within the BREX system and highlight the strength of studying inhibitors to interrogate complex systems like BREX.

## Results

### OrbA restores ICP1 genome replication in the presence of BREX

When *orbA* is deleted from ICP1 (ICP1 Δ*orbA*), ICP1 cannot form plaques in the presence of the SXT *Vch*Ind5 encoding the BREX system (BREX^Ind5^), but this phage maintains the ability to plaque equally well on the laboratory strain of *V. cholerae* lacking an SXT (permissive) and in the isogenic background where BREX has been removed from the SXT *Vch*Ind5 by deletion of the hotspot 5 region (ΔBREX^Ind5^) (Fig 1A)^25^. The type I BREX system encoded by *Bacillus cereus* was previously shown to inhibit the genome replication of the phage Φ3T when expressed in *Bacillus subtilis*^15^. To determine if BREX^Ind5^ restricts ICP1 genome replication in the absence of OrbA, we evaluated the fold change in ICP1 Δ*orbA* genome copy number in the presence and absence of BREX^Ind5^ by quantitative PCR (qPCR) over a single round of infection. ICP1 produces new progeny by 20 minutes post-infection, so time points were chosen to represent discrete phases in the ICP1 genome replication cycle: 4- (pre-replication), 8- (theta replication), 16- (rolling circle replication), and 20 minutes post-infection^31^. In the permissive and ΔBREX^Ind5^ backgrounds, ICP1 Δ*orbA* replicated robustly, while in the presence of BREX^Ind5^, ICP1 genome replication was inhibited by 8 minutes post-infection, indicating that the BREX^Ind5^ system encoded by *Vch*Ind5 inhibits ICP1 genome replication before the start of theta replication (Fig 1B).

**Figure 1.**
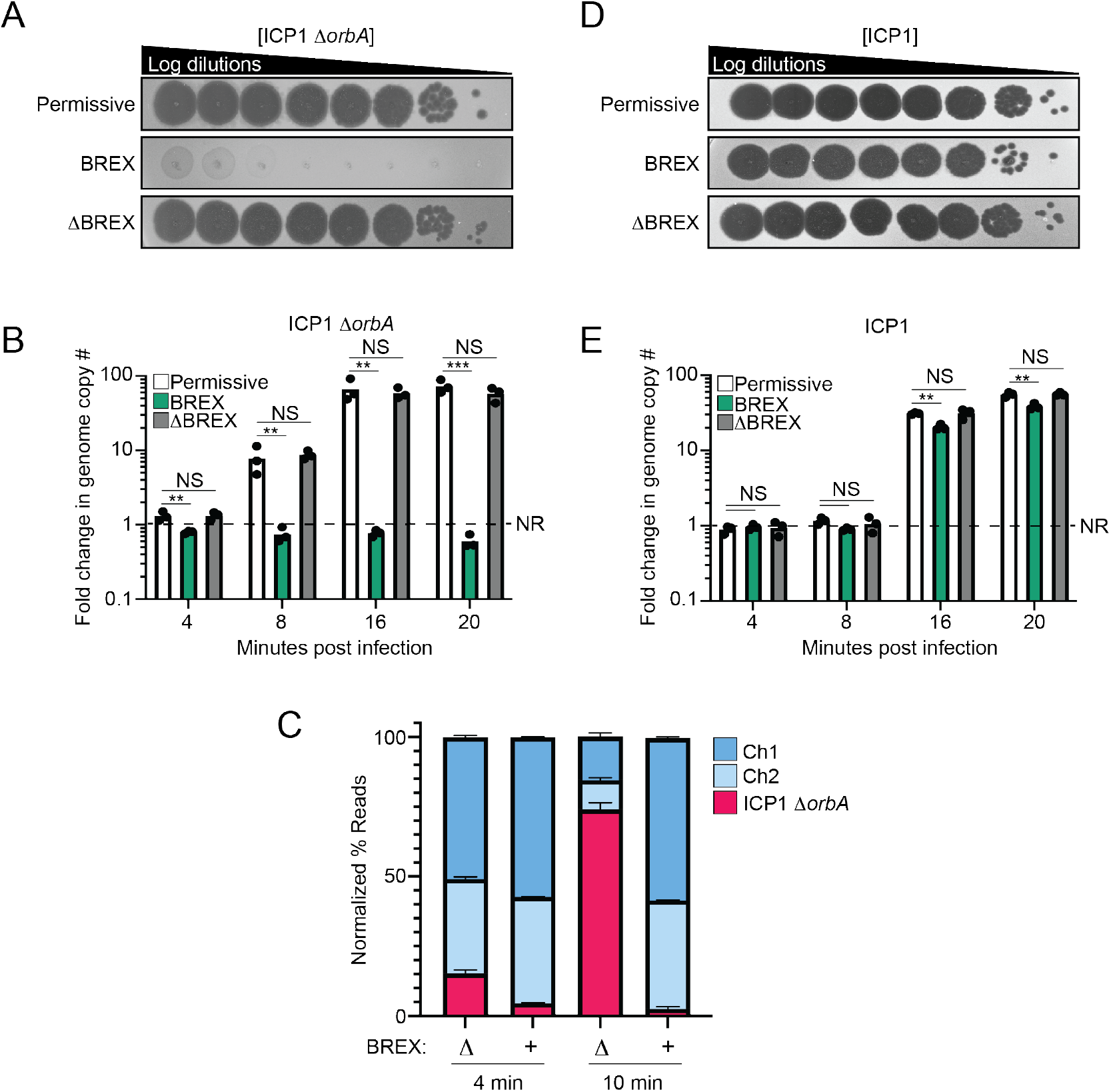
OrbA restores phage genome replication in the presence of BREX. **A)** 10-fold serial dilutions of ICP1 Δ*orbA* spotted on a lawn of bacteria (grey background). **B)** Quantification of ICP1 Δ*orbA* genome replication in the presence and absence of BREX. *V. cholerae* strains were infected at a multiplicity of infection (MOI) of 0.1. Samples were taken at the addition of phage (Time 0) and at the indicated time points. Each dot represents a biological replicate. NR - no replication. Statistical analysis was performed using a one-way ANOVA (* = p<0.01, ** = p<0.001, *** = p<0.0001, **** p=<0.00001). NS – not significant **C)** Deep sequencing of total DNA extracted from cells infected with ICP1 Δ*orbA* at an MOI 1 at the indicated time points. The percentage of reads mapping to either the *V. cholerae* chromosome 1 or 2 (Ch1 or Ch2) or ICP1 was calculated by dividing the number of reads by the length of each element and then dividing by the total number of reads of all elements. **D)** 10-fold serial dilutions of ICP1 spotted on the same lawns of bacteria (grey background) in panel A. **E)** Quantification of ICP1 genome replication in the presence and absence of BREX, as in panel B. For all panels, The *V. cholerae* strain E7946 was used as a permissive background (permissive), E7946 encoding the SXT ICE *Vch*Ind5 is indicated as BREX, and E7946 encoding the SXT ICE *Vch*Ind5 with a deletion of the entire hotspot 5 region is indicated as ΔBREX.

Inhibition of phage genome replication is generally thought to be caused by the degradation of the phage genome; however, the *B. cereus* BREX system did not degrade the Φ3T genome as determined by Southern blot analysis^15^. Bioinformatic analysis of BREX systems has not identified a nuclease domain in any BREX componenent^15^, and experimentally, lysates from cells expressing the BREX system mixed with phage DNA did not show detectable nuclease activity^16^, further supporting the observation that the BREX system does not degrade the phage genome. A limitation of our qPCR results and the study on the *B. cereus* BREX system is that both conclusions are based on using a single region of the phage genome to infer the condition of the phage genome globally^15^.To ensure that the qPCR results were representative of the phage genome as a whole, we performed a deep-sequencing experiment where we sequenced the total DNA from cells infected by ICP1 Δ*orbA* in the presence and absence of the BREX^Ind5^ system before genome replication (4 minutes) and after theta replication had begun (10 minutes). In the absence of BREX^Ind5^, and in line with our qPCR results, the number of reads mapping to the ICP1 Δ*orbA* genome was robust at 10 minutes post-infection, whereas, in the presence of BREX^Ind5^, the number of reads mapping to the ICP1 Δ*orbA* genome appeared the same at both time points, indicating that the ICP1 Δ*orbA* did not replicate (Fig 1C). We next determined the mean read coverage across the ICP1 Δ*orbA* genome to assess if coverage loss varied across the genome. We found that the mean read coverage was consistent across the genome in the presence and absence of BREX at 4 minutes post-infection (Fig S1A). At 10 minutes post-infection, in the presence of BREX, read coverage was consistent across the phage genome, whereas, in the absence of BREX, the ICP1 Δ*orbA* genome had a coverage profile indicative of theta replication (Fig S1B). Together, our results indicate that the BREX system inhibits ICP1 Δ*orbA* by preventing the phage from initiating genome replication.

Wild-type OrbA-positive ICP1 (ICP1) can form plaques in the presence of BREX^Ind5^, but the efficiency of plaquing is reduced about 4-fold when compared to the permissive or ΔBREX^Ind5^ backgrounds (Fig 1D and Fig S1C). The slight reduction in ICP1 plaquing in the presence of BREX^Ind5^ suggests that while OrbA can restore ICP1 plaque formation, OrbA may not completely inhibit the BREX^Ind5^ system. To assess if ICP1 genome replication also shows a slight defect in the presence of OrbA, we determined the fold change in ICP1 genome copy number in the presence and absence of BREX^Ind5^ by qPCR over a single round of infection. ICP1 genome replication was robust under all conditions; however, we note a slight 2-fold decrease in ICP1 replication in the presence of BREX^Ind5^ (Fig 1E), in line with our observation that there is a slight reduction in ICP1 plaquing in the presence of BREX^Ind5^ (Fig 1D and Fig S1C). Together, these results demonstrate that the Type I BREX^Ind5^ system in *V. cholerae* restricts phage genome replication and that inhibition of BREX^Ind5^ by OrbA largely restores ICP1 genome replication.

### OrbA directly binds and alters the interaction between BrxC subunits

How OrbA inhibits BREX^Ind5^ to restore ICP1 genome replication is unknown. To determine if OrbA inhibits BREX^Ind5^ by directly interacting with a BREX component or host protein, we took an unbiased approach and performed an *in vivo* co-immunoprecipitation (CoIP) using C-terminally 3XFLAG tagged OrbA (OrbA-FLAG) that retained its function (Fig S2A) and expressed better than the N-terminal version (Fig S2B). Since OrbA can protect two unrelated phages from BREX^Ind5 25^, we hypothesized that a phage component is not required for OrbA function and thus, we performed the OrbA CoIPs in the absence of phage infection. Lysates, where either OrbA-FLAG or the FLAG tag control (FLAG) were expressed in the presence of the BREX^Ind5^ system in its native context, were incubated with anti-FLAG resin and subsequently eluted using FLAG peptide. Elutions were resolved by SDS-PAGE, and faint bands were observed in the FLAG control eluate, whereas a dominant band indicative of OrbA-FLAG together with a higher molecular weight band was observed in the experimental elutions (Fig 2A and Fig S3). To identify the putative interaction partners, eluate from the OrbA-FLAG and the FLAG control were analyzed by mass spectrometry. The most abundant protein enriched in the presence of OrbA-FLAG was the ATPase BrxC (WP_000152592), encoded by the BREX^Ind5^ system (Table S1). These results suggest that BrxC is the putative interaction partner of OrbA.

**Figure 2.**
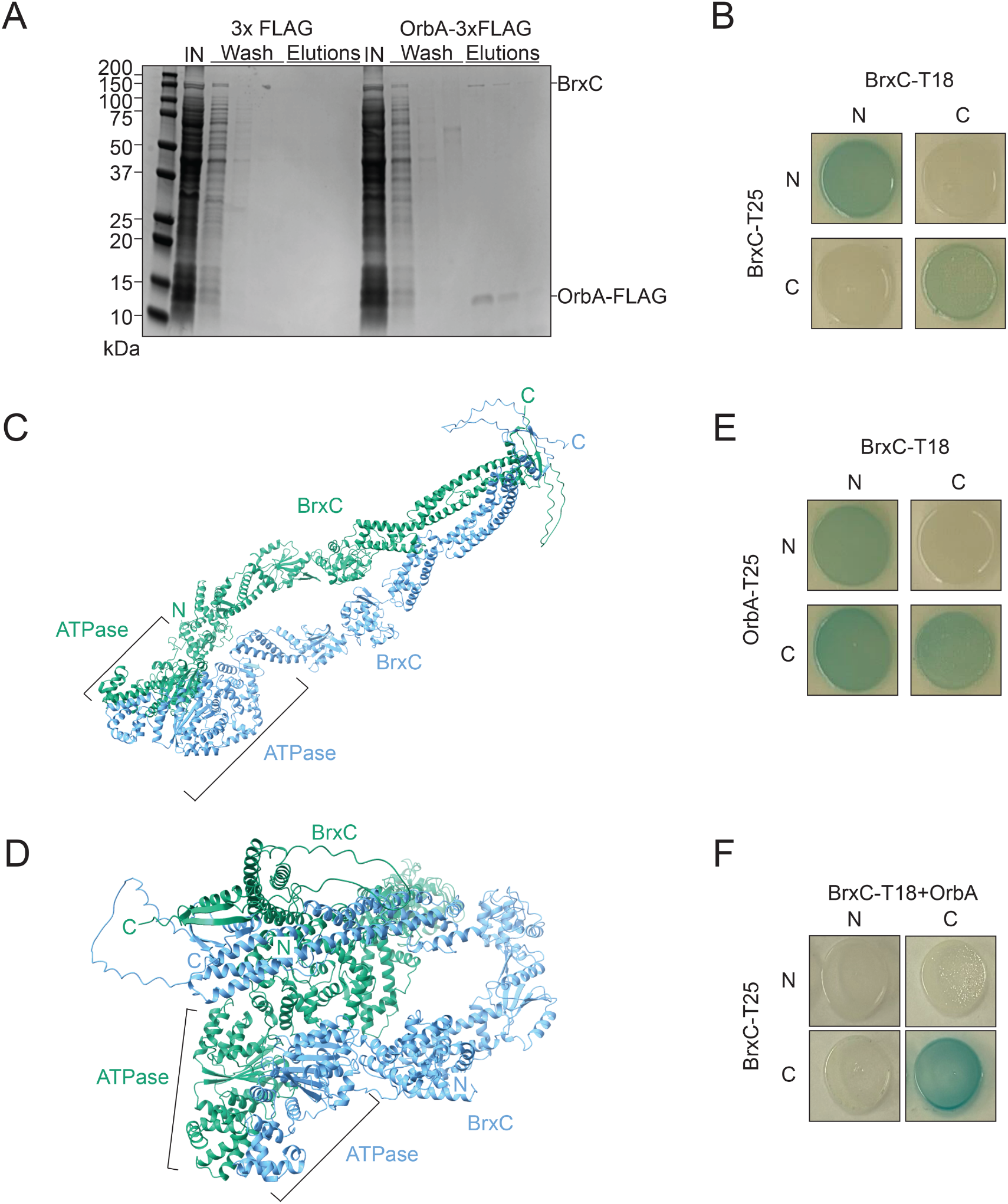
OrbA directly interacts with BrxC from *Vch*Ind5. **A)** A representative gel image of samples from 3xFLAG and OrbA-3xFLAG resolved by SDS-PAGE and stained with Coomassie brilliant blue. IN – input. For additional biological replicates, see Figure S3. **B)** Bacterial two-hybrid analysis of the interaction between BrxC-BrxC. N- and C- indicate the terminus of each half of the adenylate cyclase enzyme that was translationally fused to the protein of interest. **C-D)** Structural prediction of a BrxC dimer by ColabFold. **C)** Open conformation of the BrxC dimer. We note that the N-terminus of the right dimer is behind the ATPase and is not visible. **D)** Closed conformation of the BrxC dimer. N – N terminus. C – C terminus. **E-F)** Bacterial two-hybrid analysis of the interaction between **E)** BrxC-OrbA and **F)** BrxC-BrxC with OrbA also being expressed from the T18 plasmids. Performed the same as in panel B.

To validate the interaction between OrbA and BrxC, we performed a bacterial two-hybrid (BACTH) assay, where the N- or C- terminus of each protein of interest was translationally fused to either half (T18 or T25) of the enzyme adenylate cyclase^34^. If the protein pairs interact, colonies turn blue in the presence of X-gal (5-Bromo-4-Chloro-3-Indolyl β-D-Galactopyranoside), whereas if the pairs do not interact, colonies remain white. The combination of BrxC-BrxC, where the two halves of adenylate cyclase were present on the same termini, led to the formation of blue colonies, while the combinations where the opposite termini are tagged did not show a color change (Fig 2B). These results indicate that BrxC can interact with itself and most likely forms a multimer. To assess the BrxC-BrxC interaction, we determined the predicted structure of a BrxC dimer using ColabFold^35^. The predicted dimer of BrxC indicates that the N-terminus of one subunit interacts with the N-terminus of another, the same is true for the C-terminus (Fig 2C). Additionally, the predicted structure indicates that the N- and C-terminus of the BrxC subunits are separate from each other and at a distance (∼8Å) that would not support an interaction between the two termini. ColabFold also predicts a second conformation for the BrxC dimer where the BrxC subunits maintain their N- and C-terminal dimerization, but the subunits fold over and bring the N-termini close to the C-termini (Fig 2D). While this second conformation could be possible, our BACTH results did not capture an interaction between the N- and C- termini. Together, our results indicate that BrxC interacts with itself to form a multimer.

The combination of OrbA-OrbA resulted in white colonies, indicating that OrbA does not interact with itself (Fig S4A). Expression of either N- or C-terminally tagged OrbA in combination with N-terminally tagged BrxC resulted in blue colonies (Fig 2E and Fig S4B). All combinations of C-terminally tagged BrxC resulted in no blue colonies except for the combination of C-terminally tagged BrxC with T18 and C-terminally tagged OrbA with T25 resulted in a positive interaction (Fig 2E and Fig S4B). The difference in results with the C-terminally tagged OrbA could be the result of the tag position and its proximity to the C-terminal tag of BrxC. Together, these results support an interaction between OrbA and BrxC. We next sought to determine if OrbA binding would alter how BrxC interacts with itself by performing a modified BACTH assay. Using one plasmid from the wild-type BrxC pairings (T18), we introduced untagged OrbA with a ribosome binding site downstream of the BrxC open reading frame. Expression of OrbA with the wild-type BrxC pairing showed a positive interaction when both C-termini were tagged like that observed for the wild-type BrxC pairings, whereas the interaction observed when the N-termini were tagged was lost (Fig 2F). We conclude that OrbA binding alters how BrxC subunits interact with each other, and we predict that the altered interaction disrupts BrxC function and restores ICP1 genome replication.

### BrxC is necessary but not sufficient to inhibit phage genome replication

BrxC has been previously shown to be necessary for BREX-mediated phage defense in systems found in *E. coli* HS, *B. cereus*, and *Acinetobacter* species NEB394 when expressed in a heterologous host^15,16,19^. To confirm the necessity of BrxC under native conditions, a deletion of *brxC* in *V. cholerae* harboring the SXT ICE *Vch*Ind5 was generated, and the resulting strain was challenged with ICP1 Δ*orbA*. The efficiency of plaquing (EOP) of ICP1 Δ*orbA* on a wild-type BREX^Ind5^ background was at the limit of detection, whereas infection of the Δ*brxC* derivative restored the EOP to 1 (Fig 3A), demonstrating that BrxC is necessary for phage defense. To confirm that the abrogation of phage defense was not due to a polar effect on the downstream genes, a plasmid encoding BrxC under an inducible promoter was introduced into the *brxC*-deletion background. Induction of BrxC restored phage restriction to nearly the same degree as a wild-type BREX^Ind5^ system, whereas an empty vector control did not, demonstrating that the loss of phage defense was not due to a polar effect (Fig 3A). Furthermore, the expression of BrxC in the absence of the other BREX components failed to protect against ICP1 Δ*orbA* infection, showing that BrxC alone is insufficient for phage defense (Fig 3A). We conclude that BrxC is necessary for phage defense, but other components of the BREX^Ind5^ system are required for protection.

**Figure 3.**
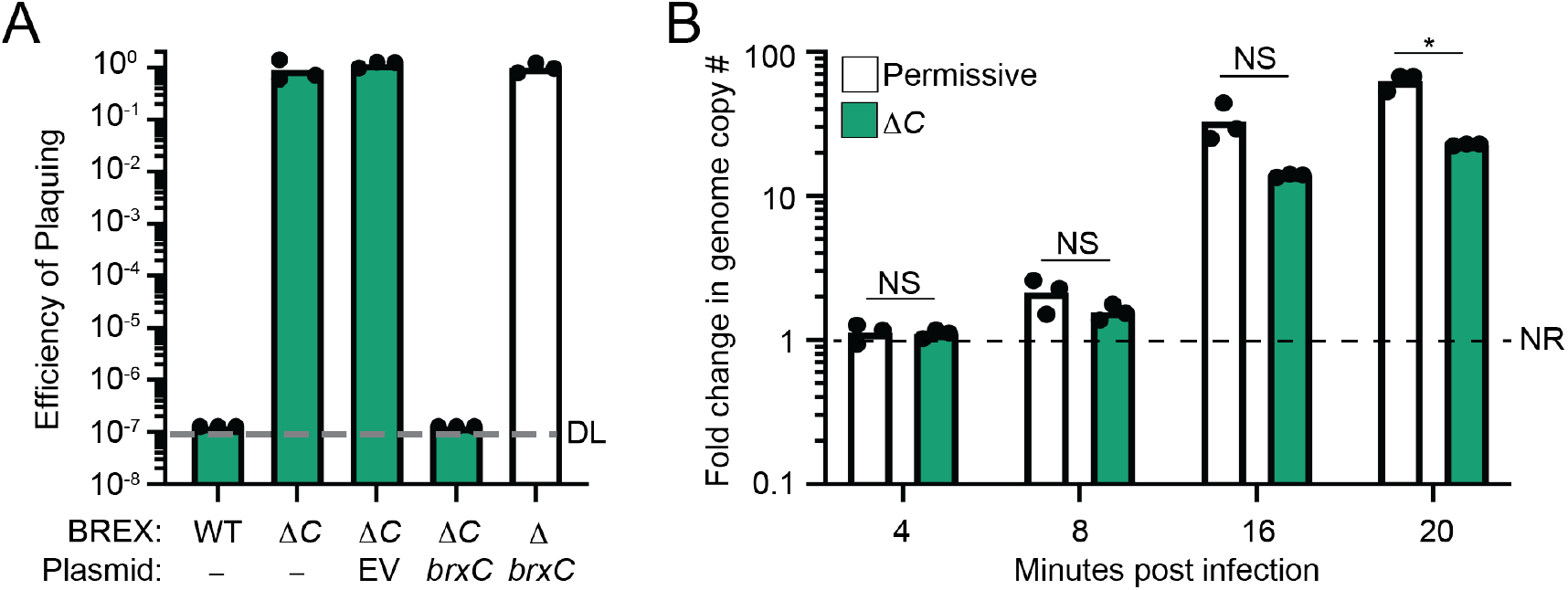
BrxC is required for inhibition of phage genome replication. **A)** Efficiency of plaquing (EOP) of ICP1 Δ*orbA* on *V. cholerae* with the genotype indicated relative to the ΔBREX strain background. ΔC – deletion of *brxC*. EV - empty vector. DL - detection limit. Each dot represents a biological replicate **B)** Quantification of ICP1 Δ*orbA* replication in a ΔBREX and *brxC*-deletion background (ΔC). NR - no replication. Statistical analysis was performed using a student t-test was performed (* = p<0.01, ** = p<0.001, *** = p<0.0001, **** p=<0.00001). NS – not significant. Each dot represents a biological replicate.

Since the BREX^Ind5^ system impairs phage genome replication and we observed a slight reduction in ICP1 replication in the presence of BREX^Ind5^ (Fig 1E), we sought to determine if the absence of BrxC would restore ICP1 genome replication to the efficiency observed when infecting a ΔBREX^Ind5^ background. To test this, we performed qPCR measuring the fold change in ICP1 Δ*orbA* genome copy number over a single round of infection in the *brxC*-deletion background. The replication of ICP1 Δ*orbA* was robust in the absence of BrxC, as expected, given the observation that ICP1 Δ*orbA* plaque formation was restored in this mutant background (Fig 3B). However, ICP1 Δ*orbA* replication was still 2-fold down in the Δ*brxC* mutant in comparison to the permissive control, phenocopying the replication observed for OrbA-proficient ICP1 in the presence of BREX^Ind5^ (Fig 1E). These results suggest that phage genome replication is restored by OrbA disrupting BrxC function in a way that phenocopies the absence of BrxC. We conclude that BrxC is required for robust inhibition of phage genome replication, but in the absence of BrxC, the BREX system retains some limited function and reduces phage genome replication.

### BrxC ATPase activity is required for BrxC-BrxC interaction and BREX-mediated defense

Structural prediction and domain architecture analysis of BrxC suggests that the N-terminus of BrxC has a putative ATPase domain that shares homology with the ATPase domain of the <u>o</u>rigin recognition <u>c</u>omplex subunit 4 (ORC4) found in both Eukaryotes and Archaea^36–39^. To determine if the predicted ATPase activity of BrxC is required for phage defense, we identified the Walker A motif - required for ATP binding, and the Walker B motif - necessary for ATP hydrolysis, and mutated key residues to Alanine to disrupt function. To identify the Walker A motif, we searched the BrxC primary sequence for a conical Walker A motif (GxxxxGKT)^40,41^. We identified one instance of this motif and mutated the key lysine residue at position 73 (Fig S5A). The Walker B motif is identified as a group of four hydrophobic residues proceeded by a dyad of aspartic and glutamic acids, with the glutamic acid being the key residue for ATP hydrolysis^41^. Together, the Walker A and B motifs form a catalytic pocket, so to identify the BrxC Walker B motif, we used the predicted BrxC ATPase (residues 1-532) dimer structure and used the lysine residue at position 73 as a marker for the location of a putative catalytic pocket (Fig S5A). We identified two glutamic acids within the putative catalytic pocket at positions 255 and 275 (Fig S5A). The glutamic acid at position 275 was not considered for further investigation as it is not paired with an aspartic acid that is important for the coordination of ATP; therefore, we predicted that the glutamic acid at position 255 is the key residue for ATP hydrolysis and mutated it to an alanine.

The wild-type *brxC* allele and the Walker A (*brxC*^K73A^) and Walker B (*brxC*^E255A^) mutants were introduced into a plasmid under an inducible promoter and translationally fused to a C-terminal 3xFLAG tag. Each construct was introduced into a *brxC*-deletion background and tested for the ability to restore phage restriction. The expression of wild-type BrxC restored inhibition of ICP1 Δ*orbA* in comparison to the empty vector control (Fig 4A). However, neither BrxC^K73A^ nor BrxC^E255A^ was sufficient to complement the deletion of *brxC* (Fig 4A). To verify that loss of complementation was not due to poor expression of the mutant alleles, we performed a Western blot using a primary antibody against the FLAG tag. Both mutant alleles were expressed at levels comparable to each other but appeared to be slightly reduced when compared to the wild-type allele (Fig S5B), so while protein abundance may have a small impact, we expect that the altered activity of these two proteins is the dominant reason for the loss of complementation. From these results, we infer that both ATP binding and hydrolysis are required for BrxC function and phage defense.

**Figure 4.**
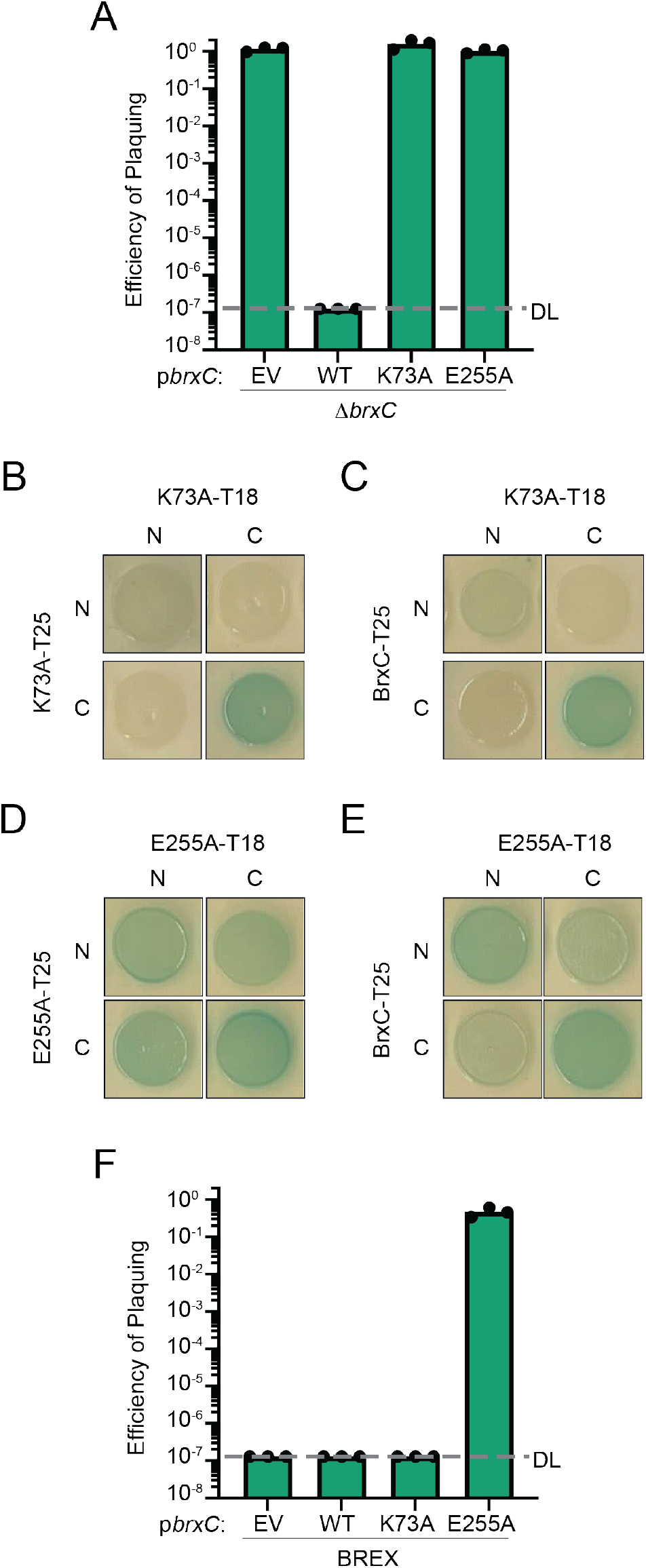
ATP-binding and hydrolysis activity of BrxC is required for function and interactions. **A)** Efficiency of plaquing (EOP) of ICP1 Δ*orbA* on a *brxC*-deletion background harboring a plasmid that either encodes wild-type BrxC, the Walker A (K73A) or Walker B (E255A) mutant relative to the *brxC*-deletion strain harboring an empty vector. DL - detection limit. EV - empty vector. Each dot represents a biological replicate. **B-E)** Bacterial two-hybrid analysis of the interaction between **B)** BrxC^K73A^-BrxC^K73A^, **C)** BrxC^K73A^-BrxC, **D)** BrxC^E255A^-BrxC^E255A^, and **E)** BrxC^E255A^-BrxC. N- and C- indicate the terminus of each half of the adenylate cyclase enzyme was fused to the protein of interest. **F)** EOP of ICP1 Δ*orbA* on *V. cholerae* encoding a wild-type BREX system and harboring a plasmid that expresses the allele of BrxC indicated relative to the permissive strain *V. cholerae* E7946 lacking *Vch*Ind5 and harboring an empty vector. Each dot represents a biological replicate.

Many ATPase-domain-containing proteins have been shown to form multimers regulated through the interaction with ATP^40,41^. For example, ORC4, which the ATPase domain of BrxC shows structural homology to, forms a heterohexameric complex with ORC subunits 1-3, 5, and 6 dependent on ATP binding^42,43^. Our BACTH results and structural prediction of the BrxC dimer (Fig 2), support the hypothesis that BrxC forms a multimer. To assess if ATP binding and hydrolysis are required for BrxC subunits to interact, we performed BACTH assays using wild-type BrxC and both the Walker A (BrxC^K73A^) and Walker B (BrxC^E255A^) mutants. While wild-type BrxC-BrxC pairing showed interactions between the N-N and C-C termini-tagged pairings (Fig 2B), the interaction between BrxC^K73A^-BrxC^K73A^, maintained an interaction when both C-termini were tagged but showed a reduced interaction when both N-termini were tagged in comparison to wild-type BrxC-BrxC interaction, suggesting that ATP binding is required for robust interaction (Fig 4B). Additionally, as in the wild-type BrxC-BrxC interactions, no interaction between the N-C termini pairings was observed. We next tested if BrxC^K73A^ could interact with a wild-type BrxC and observed an interaction pattern similar to the BrxC^K73A^-BrxC^K73A^ pairing, suggesting that efficient N-N terminal interactions require both BrxC monomers to be ATP-binding proficient (Fig 4C and S5C). Additionally, this result phenocopies the interaction pattern observed when OrbA is expressed during the BrxC-BrxC interaction (Fig 2F), suggesting that OrbA may alter BrxC multimerization by disrupting BrxC from interacting with ATP.

In the case of the interaction between two BrxC^E255A^ monomers, both wild-type BrxC-BrxC interactions were maintained, but additional interactions were observed between the N-terminal and C-terminal tagged pairings (Fig 4D). These interactions were also observed between the wild-type BrxC and the BrxC Walker B mutant (Fig 4E and S4D). The change in the interaction profile suggests that BrxC goes through a conformational change that brings the N-terminus closer to the C-terminus, and this structural change is supported by our predicted BrxC dimer that shows a closed conformation, bringing the N-terminus in proximity to the C-terminus (Fig 2D). Together, these results suggest that BrxC goes through conformational changes that are important for BrxC function. We next assessed if mutation of the Walker A or B motif altered OrbA interaction and found that both variants maintained an interaction with OrbA, suggesting that the interaction does not depend on ATP binding or hydrolysis (Fig S5E-F)

Since BrxC^E255A^ maintains the ability to interact with wild-type BrxC, we hypothesized that BrxC^E255A^ would act as a dominant-negative over the wild-type BrxC. To test this, we expressed either wild-type BrxC, the Walker A or Walker B mutant in the presence of BREX^Ind5^ and challenged each strain with ICP1 Δ*orbA*. Expression of wild-type BrxC and the Walker A mutant did not disrupt BREX^Ind5^ function as ICP1 Δ*orbA* failed to plaque on both backgrounds (Fig 4F). However, expression of the Walker B mutant *in trans* restored ICP1 Δ*orbA* plaquing, indicating that the Walker B mutant acts dominant over the wild-type BrxC and the interaction between the BrxC^E255A^ mutant and wild-type BrxC disrupts BREX^Ind5^ function (Fig 4F). We conclude that ATP binding is required for the efficient interaction between the N-terminal domains of BrxC and that BrxC, in the absence of ATP hydrolysis, is locked in an inactive state that abrogates phage defense.

### OrbA cannot inhibit the Type I BREX system encoded by SXT ICE *Vch*Ban9

In addition to the BREX system encoded by the SXT ICE *Vch*Ind5, the SXT ICE *Vch*Ban9 also encodes a Type I BREX system (BREX^Ban9^), which encodes components that share between 16-44% amino acid identity with those of BREX^Ind5^ (Fig 5A)^25^. BREX^Ban9^ robustly inhibits ICP1, indicating that OrbA is not sufficient to inhibit all Type I BREX systems^25^. Furthermore, BREX^Ban9^ inhibited ICP1 genome replication (Fig S6A) in parallel to our findings with BREX^Ind5^. The inability of OrbA to inhibit BREX^Ban9^ was unsurprising as the BrxC proteins share low amino acid identity (21%), and our results thus far support the model that OrbA disrupts BrxC^Ind5^ function through a direct interaction. However, BREX^Ban9^ encodes a Type IV restriction enzyme BrxU^18^, that could provide protection against ICP1 and complicate our interpretation (Fig 5A). However, we found that ICP1 was inhibited to the same level in the presence and absence of BrxU, indicating that phage restriction is not due to BrxU and that OrbA cannot inhibit BREX^Ban9^ (Fig S6B). Since the BrxC from BREX^Ban9^ shares low amino acid identity with BrxC from BREX^Ind5^, we hypothesized that OrbA fails to inhibit BREX^Ban9^ because OrbA cannot interact with BrxC^Ban9^. To test this hypothesis, we performed a BACTH assay. Neither combination of OrbA with BrxC^Ban9^ showed a positive interaction, supporting the hypothesis that these two proteins do not interact (Fig 5B). We conclude that OrbA is a specific inhibitor against the BREX^Ind5^ system and not a general inhibitor against all Type I BREX systems.

**Figure 5.**
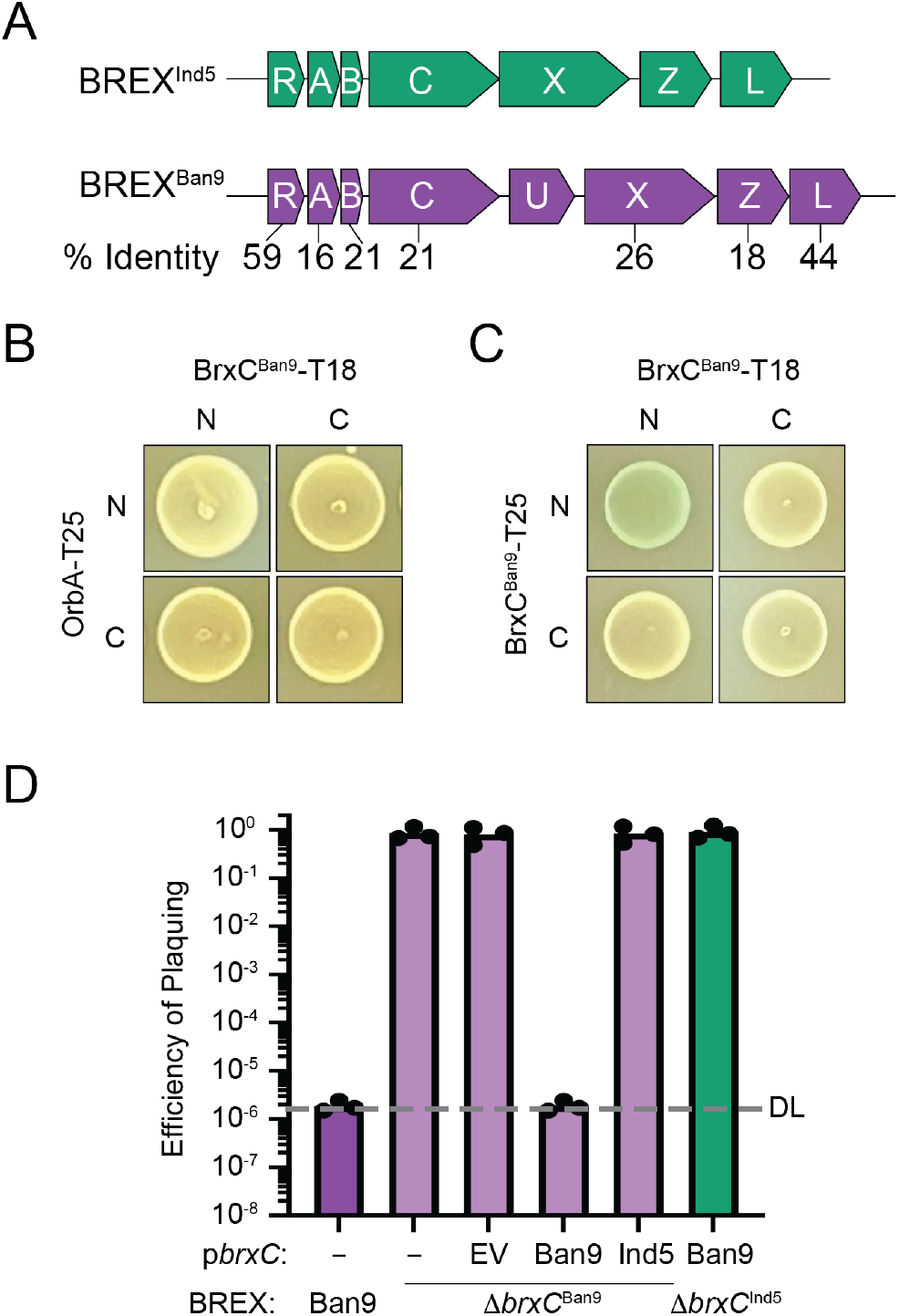
BrxC^Ban9^ does not interact with OrbA and cannot functionally complement a *brxC*^Ind5^-deletion. **A)** Schematic of the BREX system encoded by SXT ICE *Vch*Ind5 (BREX^Ind5^) and *Vch*Ban9 (BREX^Ban9^) (not drawn to scale). The percent amino acid identity between each BREX protein is indicated below each BREX component: BrxR (R), BrxA (A), BrxC (C), BrxU (U), BrxX (X), BrxZ (Z), and BrxL (L). **B-C)** Bacterial two-hybrid analysis of the interaction between **B)** BrxC^Ban9^-OrbA and **C)** BrxC^Ban9^-BrxC^Ban9^. N- and C- indicate the terminus of each half of the adenylate cyclase enzyme was fused to the protein of interest. **D)** Efficiency of plaquing (EOP) of ICP1 on *V. cholerae* either encoding a wild-type BREX system from *Vch*Ban9 (Ban9), a BREX system from *Vch*Ban9 with *brxC* deleted (Δ*brxC*^Ban9^), or a BREX system from *Vch*Ind5 with *brxC* deleted (Δ*brxC*^Ind5^) and the plasmid expressing *brxC* as indicated relative to either the permissive *V*. cholerae E7946 lacking an SXT or E7946 lacking an SXT ICE and harboring an empty vector plasmid. Each dot represents a biological replicate. DL - detection limit. EV - empty vector.

While both alleles of the BrxC protein have different sensitivity to OrbA, their predicted dimer structures by ColabFold are similar, indicating they likely have the same function (Fig S7). To validate that BrxC^Ban9^ interacts with itself, we performed a BACTH assay (Fig 5C). We observed an interaction when both N-termini were tagged but did not observe an interaction when both C-termini were tagged or when the N-terminal tag was paired with the C-terminal tag. From these results, we predict that BrxC^Ban9^ can multimerize; however, the requirement for multimerization may differ from BrxC^Ind5^. We next sought to determine if the BrxC^Ban9^ allele could functionally complement a BrxC^Ind5^ deletion and provide protection against OrbA-proficient ICP1. We first generated a deletion of BrxC^Ban9^ and challenged the deletion strain with ICP1 Δ*orbA*; in line with previous observations, a BrxC^Ban9^-deletion mutant failed to inhibit phage infection (Fig 5D). Moreover, inhibition of ICP1 *ΔorbA* was restored when BrxC^Ban9^ was expressed from a plasmid in the BrxC^Ban9^-deletion background. These results further support the importance of BrxC between diverse Type I BREX systems. To test for functional complementation, each BrxC inducible construct was introduced into both the BrxC^Ind5^- and BrxC^Ban9^-deletion mutants. The expression of each BrxC allele restored phage inhibition when in combination with their respective system, whereas an empty vector control did not (Fig 5D and 3A). The expression of the reciprocal BrxC protein in each BrxC-mutant background failed to complement, indicating that while these two proteins have similar predicted activities, the inability to functionally complement may be due to the loss of interactions between BrxC and other components (Fig 5D). We conclude that each BrxC has evolved to function with its respective BREX components.

### Homologs of BrxC^Ind5^ are more diverse than BrxC^Ban9^

OrbA encoded by ICP1 does not interact directly with BrxC^Ban9^, so it fails to inhibit the BREX^Ban9^ system. We sought to identify OrbA homologs that may be able to inhibit BREX^Ban9^ using PSI-BLAST. From this analysis, we did not identify any OrbA homologs outside of ICP1. To potentially address why OrbA cannot inhibit BrxC^Ban9^, we next generated a phylogenetic tree of BrxC homologs within the NCBI-nr database using BrxC^Ind5^ and BrxC^Ban9^ as queries for PSI-BLAST searches. We limited the results to 1860 distinct homologs that grouped into eight clades, with BrxC^Ind5^ and BrxC^Ban9^ separating into different clades, clade V and clade III, respectively (Fig 6). Along with BrxC^Ban9^, BrxC proteins from previously characterized BREX systems from *E. fergusonii, E. coli HS, S. enterica*, and *Acinetobacter* spp also separated into clade III while the BrxC protein characterized from *B. cereus* is in clade II (Fig 6). BrxC proteins from clades I – III predominantly followed the phyletic grouping of their host, whereas clades IV – VIII with BrxC^Ind5^ did not (Fig 6; inner ring). Instead, BrxC proteins in clades IV – VIII generally clustered into groups that followed their bacterial class, with BrxC^Ind5^ being part of a gammaproteobacteria cluster within clade V (Fig 6; outer ring). Within each of the clades I – IV BrxC proteins are more similar to each other, compared to clades V-VIII where BrxC is more diverse within each clade. From these results, we conclude that BrxC proteins from clade IV – VIII, including BrxC^Ind5^, have gone through more instances of diversification in comparison to the more conserved BrxC^Ban9^ and BrxC proteins in clades I - III. We infer that the diversification of BrxC proteins can afford resistance to phage-encoded inhibitors like OrbA. Additionally, with BrxC being an essential component, we predict other inhibitors function through BrxC.

**Figure 6.**
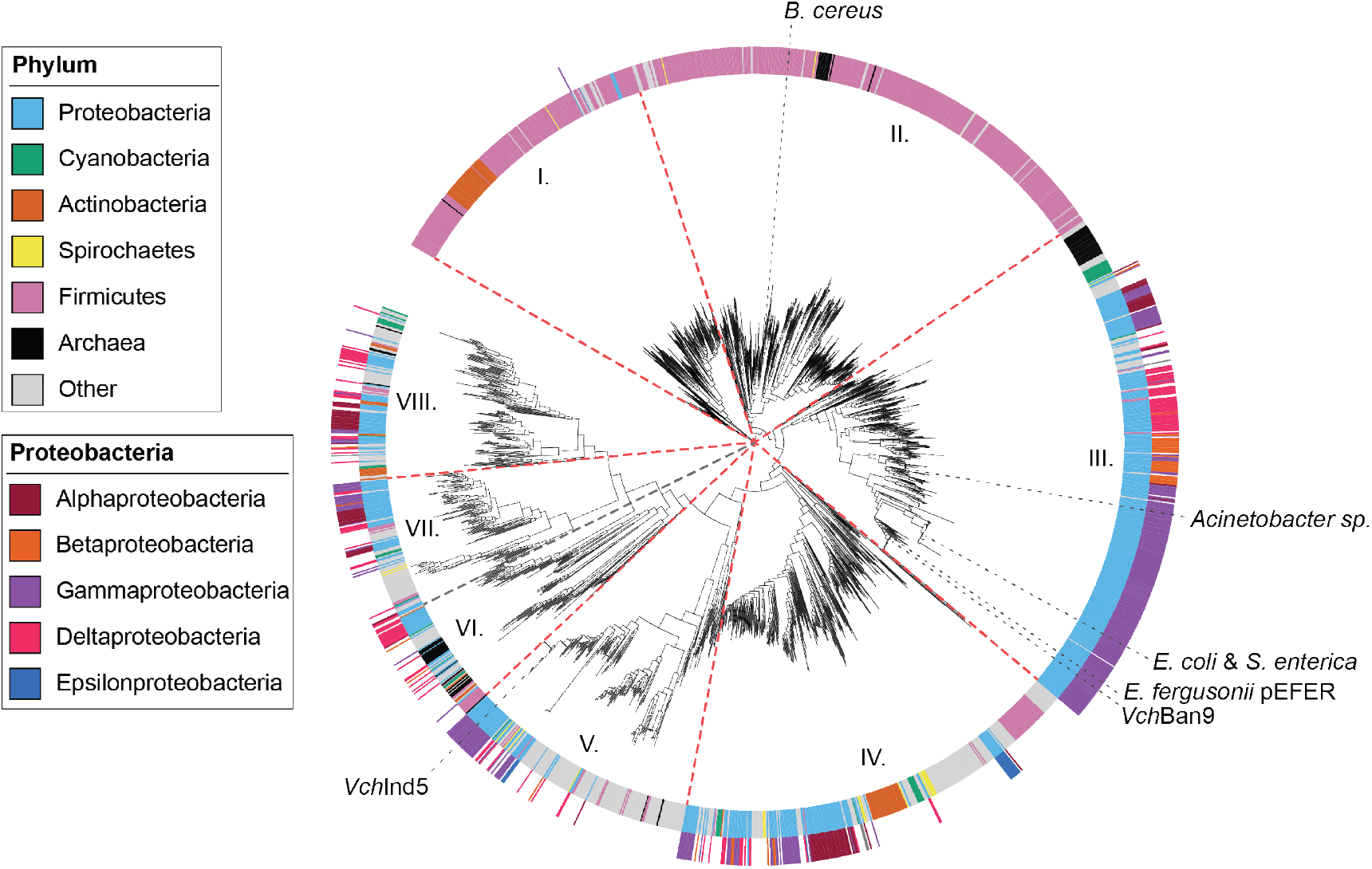
Homologs of BrxC^Ind5^ are more diverse than homologs of BrxC^Ban9^. A phylogenetic tree of BrxC homologs was generated by performing a PSI-BLAST search using the primary sequence for BrxC^Ind5^ and BrxC^Ban9^. From the search, 10,000 sequences were identified (5,000 for each allele of BrxC) that were reduced to 1,860 unique sequences sharing less than 80% amino acid identity. The inner ring indicates the phylum of the organism that encodes the allele of BrxC. The outer ring indicates the specific class of proteobacteria that encodes the allele of BrxC. The red dashed lines separate the BrxC homologs into eight different clades. The gray dashed lines indicate the alleles of BrxC from this study and the alleles of BrxC from BREX systems that have been studied elsewhere. *Bacillus cereus* (WP._001993018.1)^15^, *Acinetobacter sp*. NEB394 (WP_176538601)^19^, *Escherichia coli HS* (ABV04726.1)^16^, *Salmonella enterica* LT2 (NP_463355.1)^20^, and *Escherichia fergusonii* pEFER (WP_000687848.1)^18^.

## Discussion

Phages have evolved a diverse array of counter strategies to circumvent phage defense systems. Here, we describe how the BREX-inhibitor OrbA encoded by the vibriophage ICP1 inhibits the type I BREX system encoded by the SXT ICE *Vch*Ind5. Unlike the previously described BREX inhibitors, Ocr and SAM lyase, that inhibit the BREX system through the DNA methylase BrxX^23^ or by reducing the presence of S-adenosyl methionine (SAM)^24^, respectively, we found that OrbA inhibits BREX^Ind5^ by directly binding the ATPase BrxC (Fig 2 and 7). Together, the known BREX inhibitors use three distinct mechanisms to inhibit BREX, demonstrating that there are multiple strategies to inhibit BREX activity. The idea that there are various mechanisms to abrogate BREX function is further supported by the observation that most BREX components are required for phage restriction^15,16,19^, making each component a potential target for a phage-encoded inhibitor.

**Figure 7.**
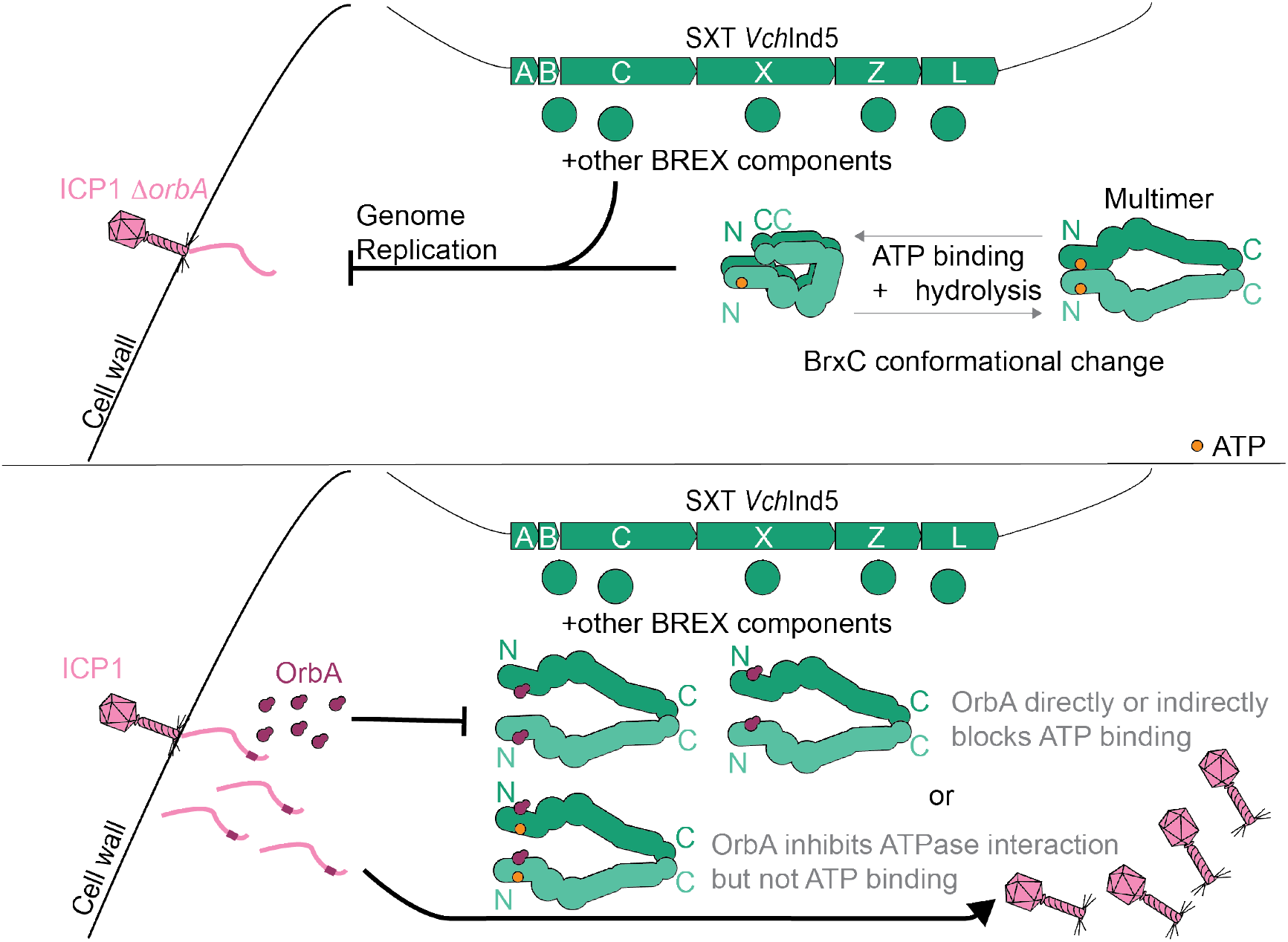
Model for BrxC conformational changes and OrbA inhibition. **Top)** *V. cholerae* infected by ICP1 Δ*orbA* in the presence of BREX that inhibits phage progeny production. BrxC forms a multimer and goes through conformational changes that are dependent on ATP binding and hydrolysis. BrxC requires other BREX components to inhibit ICP1 Δ*orbA* genome replication. **Bottom)** *V. cholerae* infected by ICP1 in the presence of BREX cannot inhibit the production of phage progeny. ICP1 expresses OrbA, and OrbA directly binds to BrxC, altering the interaction state of BrxC and disrupting its function. OrbA either inhibits ATP binding by BrxC as a direct inhibitor blocking the ATP binding site or noncompetitively by binding a different location of BrxC that disrupts ATP binding. An alternative hypothesis is OrbA does not disrupt BrxC ATP binding but prevents BrxC from multimerizing.

Ocr and SAM lyase represent broad inhibitors as they both also inhibit R-M systems^26–29^, suggesting that other inhibitors against R-M systems may also function against BREX. Current data indicates that OrbA inhibition is specific to BREX^Ind5^ as OrbA cannot inhibit BREX^Ban9^ as it does not interact with BrxC^Ban9^ (Fig 5 and Fig S6). Our phylogenetic analysis of the BrxC proteins suggests that BrxC^Ind5^ homologs (Clade V) are more diverse than the BrxC^Ban9^ homologs (Clade III) (Fig 6). We hypothesize that due to the diversity of the BrxC^Ind5^ clade as well as clades VI-VIII, inhibitors of these BrxC proteins are just as diverse and are probably specific to their BrxC, like OrbA. In comparison, BrxC^Ban9^ homologs (Clade III) show less diversification so we suspect that inhibitors against BrxC^Ban9^ homologs will be less diverse and would function against a broader range of BrxC^Ban9^ homologs. Since BREX is a common defense system, many unknown inhibitors are likely to be discovered.

While our understanding of the precise mechanism of how BREX inhibits phage genome replication remains incomplete, Ocr^23^, SAM lyase^24^, and now OrbA have helped with our understanding of how BREX functions. Ocr inhibits BREX function by acting as a DNA mimic that directly binds to BrxX, indicating that the ability of BrxX to bind to DNA is an important activity for BREX-mediated defense^23^. SAM lyase acts indirectly by reducing the availability of the substrate SAM by degrading it and inhibiting its synthesis by binding to MetK^24^. SAM is an important co-factor for methyl transferases like BrxX, suggesting that the reduction in SAM prevents BrxX from modifying the host chromosome^24,44^. Additionally, the reduction in SAM and loss of phage inhibition also suggests that SAM is important for BREX-mediated defense^24^. OrbA inhibits BREX^Ind5^ activity by directly binding to BrxC and likely disrupts how BrxC multimerizes (Fig 2). Specifically, we observed a loss of interaction between the N-termini of BrxC (Fig 2), which we infer to mean that in the presence of OrbA, the N-terminal ATPase domain of BrxC subunits cannot interact. The functional importance of BrxC ATPase domains interacting is further supported as the mutation of the Walker A motif, which is required for ATP binding, showed a reduction in N-terminal interactions and failed to complement a *brxC*-deletion, indicating that a loss of ATPase domain interaction disrupts BrxC function (Fig 4).

While mutation of the Walker A motif disrupts the interaction between ATPase domains, mutation of the Walker B motif, which is required for ATP hydrolysis, showed an altered interaction state. For the Walker B mutant, we observed interactions between the N- and C- terminus of BrxC, which is in line with the closed conformation predicted by ColabFold (Fig 2D). These results suggest that BrxC function is dependent on conformational changes, where BrxC goes from an open to a closed complex involving the N- and C-termini of BrxC subunits (Fig 4 and Fig 7). We propose a model where the ATPase domains are brought together by binding ATP, leading to a conformational change that brings the N- and C- termini near each other, and then ATP hydrolysis resets the BrxC multimer to the open conformation (Fig 7). However, in the presence of OrbA, the BrxC ATPase domains cannot interact, and thus, OrbA disrupts the function of BrxC (Fig 7). How OrbA disrupts the interaction between the ATPase domains still needs to be determined. One possibility is that OrbA directly inhibits the ability of BrxC to bind to ATP. Another possibility is that OrbA binds non-competitively and inhibits BrxC ATP-binding through an allosteric mechanism. An additional hypothesis is that OrbA does not alter the ATP binding ability of BrxC but disrupts the interaction between the ATPase domains through a mechanism independent of ATP binding (Fig 7). Together, we propose that multimerization of BrxC is required for BREX function, and OrbA inhibits BREX by altering how BrxC multimerizes.

BrxC is essential for BREX-mediated defense, and like BrxX, we demonstrate that BrxC alone is not sufficient for phage defense, providing support to the idea that BREX acts as a complex (Fig 3A). In *Salmonella enterica*, the N-terminal region of BrxC has been implicated as being important for DNA modification, while the C-terminus is important for phage restriction^20^. We can imagine a scenario where each terminus of BrxC interacts with a specific BREX component, and through the conformational change in BrxC, the activity of these other components could be regulated. However, there could be functional differences between BrxC from *S. enterica* and *BrxC* from *Vch*Ind5, as they both belong to two different clades. To better understand the mechanism of BREX inhibition, determining if BrxC interacts with other BREX components is required. Additionally, verification of the BrxC conformational changes and their function will require in-depth biochemical and structural studies. Together, our results demonstrate a new mechanism for inhibiting BREX. Through the interrogation of OrbA function, we assessed the function of the ATPase BrxC, and provided evidence that BrxC forms a multimer and goes through conformational changes. Inhibitors or phage defense systems are powerful molecular tools that will continue to provide insight into how many of these phage defense systems function. Further screening for inhibitors of BREX may provide the field with more avenues to interrogate how BREX restricts phage genome replication without degrading the phage chromosome.

## Materials and methods

### Bacterial strains and growth conditions

All *V. cholerae* and *E. coli* strains were either grown in lysogeny broth (LB) with aeration or on LB plates fortified with 1.5% agar (Fisher BioReagents). *V. cholerae* strains are derived from the strain E7946. Antibiotics, when used, were added at the following concentrations: 100 μg/mL streptomycin, 75 μg/ml Kanamycin, 100 μg/mL spectinomycin, and 1.25 or 2.5 μg/mL chloramphenicol for plasmid maintenance in *V. cholerae* when grown in liquid or on a plate respectively or 25 μg/mL chloramphenicol when growing *E. coli*. When inducer was required, either 1 mM β-D-thiogalactopyranoside (IPTG) with 1.5 mM theophylline (induction conditions for OrbA) or 20 μM IPTG with 30 μM theophylline (induction conditions for BrxC and derivatives) was used. Cells were induced either at the time of back dilution (BrxC and derivatives) or at an OD ∼0.2, and cells were grown for an additional 20 minutes at 37°C (OrbA) and then maintained throughout all other growth conditions. The genotypes of all strains used in this study are included in Table S2.

### Phage spot dilution and plaque assay conditions

Phages used in this study are listed in Table S2. Phage stocks were stored in STE buffer (100 mM NaCl, 10 mM Tris-Cl pH 8.0, and 1 mM EDTA) at 4°C. For spot dilutions, *V. cholerae* strains were grown to OD 0.3 and mixed with molten top agar (LB with 0.5% agar), overlayed on an LB plate fortified with 1.5% agar, and left at room temperature to solidify. 3μL of the appropriate phage that was 10-fold serial diluted was spotted on the top agar overlap, left to dry at room temperature, and then placed at 37°C overnight. Plaque assays used to calculate the efficiency of plating were performed by growing cells to OD 0.3, mixing 50 μL of cells with 10 μL of serially diluted phage in LB, incubated at room temperature for ∼ 10 min to allow for phage adsorption, and then each dilution was added to molten 0.5% top agar in a single petri plate. Plaque assays were incubated at 37°C overnight and then enumerated the following day. EOP was calculated as a ratio of the number of plaques on the experimental conditions over the number of plaques calculated for the permissive condition (E7946). When necessary, 2.5 μg/mL chloramphenicol, 1 mM or 20 μM IPTG, and 1.5 mM or 30 μM theophylline were added to molten 0.5% top agar to maintain both the plasmid and induction conditions.

### Generation of mutant strains and plasmid constructs

Deletions in the *V. cholerae* chromosome were generated by amplifying 1 kb arms of homology by PCR and a spectinomycin antibiotic resistance cassette flanked by FRT recognition sites (spec cassette). The arms of homology along with the spec cassette were stitched together using splicing by overlap extension (SOE) PCR and then introduced to *V. cholerae* by natural transformation^45^. *V. cholerae* cells were made naturally competent by growing the cells in LB at 30°C until an OD ∼0.3 – 1.0, pelleting the cells at 5000 x *g* for 3 minutes, washing the cells in 0.5X instant ocean, then cells were added to chitin (Sigma) slurry in 0.5X instant ocean and incubated for 24 hours at 30°C. The spec cassette was removed by the introduction of a plasmid that encodes the recombinase Flippase under an IPTG-inducible promoter and a theophylline-inducible riboswitch (riboE) by mating *V. cholerae* with *E. coli* S17 and growing the mating on 100 μg/mL streptomycin (to select for *V. cholerae*) and 2.5 μg/mL chloramphenicol (to select for the plasmid). Removal of the spec cassette and loss of the plasmid were verified by selective plating on LB plates supplemented with streptomycin, spectinomycin, or chloramphenicol. Colonies that failed to grow on both spectinomycin and chloramphenicol were chosen, and Sanger sequenced to verify that there were no mutations introduced during the recombination event.

All plasmids used in this study are listed in Table S3. Genes of interest were amplified, and plasmid linearization was accomplished by PCR. Point mutations in *brxC* were generated by amplifying *brxC* in two fragments so that the overlapping region of the up and down fragment would encode the mutation, and then two fragments were stitched together by SOE PCR. Gibson assembly was used to generate each construct. Plasmids were transformed into chemically competent *E. coli* XL-Blue cells and then verified by either Sanger sequencing or nanopore long-read sequencing at the UC Berkeley DNA sequencing facility. Verified constructs were transformed into electrocompetent *E. coli* S17 cells that are used to conjugate plasmids into *V. cholerae*. Acquisition of plasmids into *V. cholerae* was selected by plating on 100 μg/mL streptomycin and 2.5 μg/mL chloramphenicol.

### Phage genome replication by quantitative polymerase chain reaction

*V. cholerae* strains were grown to OD 0.3 and infected with either ICP1 or ICP1 Δ*orbA* strain at a multiplicity of infection of 0.1 and vortexed. Immediately, a 100 μL sample was removed and served as the zero-minute time point and boiled at 95°C for 10 minutes. Cultures were incubated at 37°C with aeration, and 100 μL samples were removed at the indicated time points, then boiled at 95°C for 10 minutes. A standard of purified ICP1 genome was made by 10-fold dilutions, while boiled samples were diluted 50-fold to serve as a template for qPCR. ICP1 genome replication was measured with primers Zac68(CTGAATCGCCCTACCCGTAC)/69 (GTGAACCAACCTTTGTCGCC) using iQ SYBR Green Supermix (Bio-Rad) and the CFX Connect RealTime PCR Detection System (Bio-Rad). Two technical duplicates were used for each biological replicate. Fold change in genome copy number was determined by dividing the calculated sq value of each time point by the zero-minute time point.

### DNA extraction, deep sequencing analysis, and read coverage

*V. cholerae* strains were grown to OD 0.3 at 37°C at 220 rpm. Cells were then infected with either ICP1 or ICP1 Δ*orbA* at a multiplicity of infection of 1 and incubated at 37°C at 220 rpm. Two milliliters of sample were collected at 4- and 10-minutes post-infection and mixed with an equal volume of ice-cold methanol. Cells were pelleted at 5000 x *g* for 5 minutes at 4°C, the supernatant was removed and resuspended in 1 mL of ice-cold 1x phosphate buffer saline pH 7.4. Cells were pelleted at 5000 x *g* for 5 minutes at 4°C, the supernatant was removed, and then pellets were stored at -80°C. Total DNA was extracted from each sample using QIAGEN DNeasy Blood and Tissue Kit and illumina sequencing was performed by Microbial Genome Sequencing Center and SeqCenter (Pittsburgh, PA) at a 200 Mb depth. The normalized percentage read coverage was calculated by taking the total number of mapped reads to each element (*V. cholerae* chromosome 1, chromosome 2, and ICP1), dividing each total by element length size to calculate reads per kilobase pair (reads/kbp), summing the total number of all three to get total number of reads per kilobase pair, and then getting the percentage by dividing the reads/kbp for each element by the total reads/kbp then multiplying by 100. The reported normalized percentage read coverage is an average of three biological replicates. The sequencing data generated for this work will be deposited in the Sequence Read Archive database and the project number will be updated here when available. Reads were mapped to a reference sequence and the mean read coverage was calculated from three biological replicates.

### Co-immunoprecipitation and mass spectrometry

*V. cholerae* strains were grown at 37°C at 220 rpm to an OD ∼0.2 in LB broth supplemented with 1.25 μg/mL chloramphenicol for plasmid maintenance then OrbA-3xFLAG and the 3xFLAG control was induced by the addition of 1 mM IPTG and 1.5 mM theophylline and grown for an additional 20 minutes. Cell pellets were collected by centrifugation at 5000 x *g* at 4°C and then stored at -80°C. Pellets were resuspended in lysis buffer (50mM Tris pH 8.0, 150 mM NaCl, 1 mM EDTA pH 8.0, 10% glycerol, 0.5% Triton X-100, and pierce protease inhibitor tablet (Thermofisher Scientific)), sonicated for 5 minutes at 20% amplitude, and cell debris was removed by centrifugation. Lysates were incubated with magnetic anti-FLAG resin (sigma) that was prewashed in Wash buffer 1 (50 mM Tris pH 8.0, 150 mM NaCl, 1 mM EDTA pH 8.0, 10% glycerol, 0.05% Triton X-100, pierce protease inhibitor tablet (Thermofisher Scientific)) and rotated by a labquake at 4°C for 2 hours. Using a magnetic stand, the supernatant was removed, and the resin was washed 3 times, once with Wash buffer 1, and twice with Wash buffer 2 (50 mM Tris pH 8.0, 150 mM NaCl,10% glycerol, and pierce protease inhibitor tablet (Thermofisher Scientific). Bound proteins were eluted off three times by incubation with 1x FLAG peptide (450ng/uL; Genscript) in 50 μL Wash buffer 2, rotated on a labquake for 30 minutes at 4°C, and then the eluate was collected. Nine microliters of input, washes, and elutions were mixed with 4x Laemmli sample buffer (Bio-Rad), boiled, and resolved on a Mini-PROTEAN TGX Precast gel (Bio-Rad), and stained with Coomassie brilliant blue. Eluates were combined and proteins were precipitated in trichloroacetic acid at a final concentration of 20% overnight at 4°C. The precipitants were centrifuged at 23,000 x *g* at 4°C for 10 minutes, the supernatant removed, and then washed three times with 1mL of 0.01 M HCl and 90% acetone. Pellets were allowed to air dry and then submitted to the Vincent J. Coates Proteomics/Mass spectrometry laboratory on the University of California – Berkeley campus for enzymatic digestion and mass spectrometry analysis.

### Western blot analysis

*V. cholerae* cells were grown to OD 0.3 as stated above in growth conditions. One milliliter of sample was spun down at 5000 x *g* for 3 minutes. Pellets were resuspended to OD 10 in 1X Laemmeli Buffer supplemented with 2-mercaptoethanol (10%) (Bio-Rad) and boiled at 99°C for 10 minutes. Ten microliters of sample were run on an Any-Kd Mini-PROTEAN TGX Precast gel (Bio-Rad) and then transferred to a Mini-size PVDF membrane (Bio-Rad) using a Trans-Blot Turbo system (Bio-Rad). Proteins were detected using a primary α-Flag antibody (Sigma) at a 1:3000 dilution and a secondary goat α-rabbit-HRP antibody (Bio-Rad) at a 1:10000 dilution. Blots were developed using Clarity Western ECL substrate (Bio-rad) and imaged using a Chemidoc XRS imaging system (Bio-Rad).

### Bacterial two-hybrid assay

One microliter of each bacterial two-hybrid plasmid was electroporated into electrocompetent *E. coli* BTH101 cells, recovered in super optimal broth with catabolite repression (SOC) for 1 hour at 37°C, then plated on LB agar plates supplemented with 50 μg/ml carbenicillin and 75 μg/ml kanamycin and incubated overnight at 37°C. Individual colonies were picked into 100μL LB broth supplemented with 50 μg/ml carbenicillin and 75 μg/ml kanamycin grown in a 96-well plate at 37°C at 200 rpm for ∼10hrs. Ten microliters of cells were spotted onto LB agar plates supplemented with 1 mM IPTG, 40 μg/mL X-Gal (5-bromo-4-chloro-3-indolyl-beta-D-galactopyranoside), 50 μg/ml carbenicillin and 75 μg/ml kanamycin, and incubated at 30 °C for 18 hours prior to imaging.

### Predicted domain and protein structure

The primary sequences of BrxC from VchInd5 was submitted to the phyre2^39^. Additionally, the primary sequence of the BrxC proteins from *Vch*Ind5 and *Vch*Ban9 were submitted to alphafold2^46^ and Colabfold^35^ using the COSMIC^2^ platform^47,48^ and were visualized using Chimera X^49^. Protein structure comparison was made using Chimera X.

### Phylogenetic tree of BrxC

Candidate proteins for BrxC phylogenetic trees were identified by iterative PSI-BLAST, using SXT ICE *Vch*Ban9 (WP_000687848) and *Vch*Ind5 BrxC (WP_000152592) proteins as queries against the NCBI nr database (accessed October 2022). Hits shorter than 1200 amino acids were removed from the database and the remaining proteins were clustered at 80% identity using CD-HIT v4.8.1^50^ to generate representative sequences throughout the tree. Representative sequences were aligned with MAFFT v7.520^51^. Phylogenetic trees were generated from the resulting alignment using FastTree 2 with default settings^51^. The resulting tree files were visualized using the iTOLv6 web browser^52^ and annotated according to annotations on the original genbank files retrieved from NCBI.

## Acknowledgments

We thank past and present Seed lab members for insightful discussion and feedback on this project. We thank Dr. Andrew Van Alst for assistance with statistical analysis. This project described was supported by Grant number R01AI153303 to K.D.S from the National Institute of Allergy and Infectious Diseases. Its contents are solely the responsibility of the authors and do not necessarily represent the official views of the National Institute of Allergy and Infectious Diseases or NIH. K.D.S. holds an Investigators in the Pathogenesis of Infectious Disease Award from the Burroughs Wellcome Fund.

