## Supplementary material for "The vibriophage-encoded inhibitor OrbA abrogates BREX-mediated defense through the ATPase BrxC": Table S2-S3 and Figures S1-S7

**Table S2: Strains**

| Strain | Genotype[a] | Reference |
|---|---|---|
| KDS6 | V. cholerae E7946, SXT(-), *Sm*$^R$, *Tm*$^S$ (O1, El Tor biotype) (referred to as permissive in text) | Laboratory collection |
| KL560 | V. cholerae E7946 Δ*lacZ::kan*, ICEVchInd5 ΔHS5::frt, *Sm*$^R$, *Tm*$^R$, *Kan*$^R$ (referred to as ΔBREX in text) | [1] |
| KS1876 | V. cholerae E7946, SXT(-) pKL06:empty vector *Sm*$^R$, *Cm*$^R$, *Tm*$^S$ | [1] |
| KS2461 | V. cholerae E7946 Δ*lacZ::kan*, ICEVchInd5 ΔfloR::frt, *Sm*$^R$, *Tm*$^R$, *Kan*$^R$ (referred to as BREX in text) | [1] |
| KS2471 | V. cholerae E7946 Δ*lacZ::kan*, ICEVchInd5 ΔfloR::frt, pGp25 (vector is pKL06::*orbA*), *Sm*$^R$, *Tm*$^R$, *Kan*$^R$,*Cm*$^R$ | [1] |
| KS2490 | V. cholerae E7946 Δ*lacZ::kan*, ICEVchInd5 ΔfloR::frt, pKL06:empty vector; *Sm*$^R$, *Tm*$^R$, *Kan*$^R$,*Cm*$^R$ | [1] |
| RO24 | V. cholerae E7946 Δ*lacZ::kan*, ICEVchInd5 ΔfloR::frt pRO01 *Sm*$^R$, *Tm*$^R$, *Kan*$^R$,*Cm*$^R$ | This study |
| RO25 | V. cholerae E7946 Δ*lacZ::kan*, ICEVchInd5 ΔfloR::frt pRO02 cm, SmR, TmR, KanR | This study |
| RO31 | V. cholerae E7946 Δ*lacZ::kan*, ICEVchInd5 ΔfloR::frt pCMB17 *Sm*$^R$, *Tm*$^R$, *Kan*$^R$,*Cm*$^R$ | This study |
| RO32 | V. cholerae E7946 Δ*lacZ::kan*, ICEVchInd5 ΔfloR (clean deletion, no frt scar) SmR, TmR, KanR | This study |
| RO38 | *V. cholerae* E7949 Δ*lacZ::kan*, ICE*Vch*Ban9 ΔfloR *Sm*$^R$, *Tm*$^R$, *Kan*$^R$, | This study |
| RO46-47 | V. cholerae E7946 Δ*lacZ::kan*, ICEVchInd5 Δ*brxC::frt-spec* ΔfloR *Sm*$^R$, *Tm*$^R$, *Kan*$^R$,*Spec*$^R$ | This study |
| RO50-51 | V. cholerae E7946 Δ*lacZ::kan*, ICEVchInd5 Δ*brxC*::*frt* ΔfloR *Sm*$^R$, *Tm*$^R$, *Kan*$^R$ | This study |
| RO53 | V. cholerae E7946 ΔlacZ::Kan, ICEVchInd5 Δ*brxC*::*frt* ΔfloR pRO11 *Sm*$^R$, *Tm*$^R$, *Kan*$^R$, Cm$^R$ | This study |
| RO54 | V. cholerae E7946 Δ*lacZ::kan*, ICEVchInd5 Δ*brxC*::*frt* ΔfloR pRO24 *Sm*$^R$, *Tm*$^R$, *Kan*$^R$, Cm$^R$ | This study |
| RO56 | V. cholerae E7946 Δ*lacZ::kan*, ICEVchInd5 Δ*brxC*::*frt* ΔfloR pKL06:empty vector cm, SmR, TmR, KanR | This study |
| RO152 | V. cholerae E7946 Δ*lacZ::kan*, ICEVchInd5 Δ*brxC*::*frt* ΔfloR pRO22 *Sm*$^R$, *Tm*$^R$, *Kan*$^R$, Cm$^R$ | This study |
| RO153 | V. cholerae E7946 Δ*lacZ::kan*, ICEVchInd5 ΔfloR pRO24 *Sm*$^R$, *Tm*$^R$, *Kan*$^R$, Cm$^R$ | This study |
| RO177 | V. cholerae E7946 Δ*lacZ::kan*, ICEVchInd5 Δ*brxC*::*frt* ΔfloR pCMB17 *Sm*$^R$, *Tm*$^R$, *Kan*$^R$, Cm$^R$ | This study |

| | | |
|---|---|---|
| RO234 | V. cholerae E7946 Δ*lacZ::kan*, ICEVchInd5 Δ*brxC*::*frt* ΔfloR pRO76 $Sm^R$, $Tm^R$, $Kan^R$, $Cm^R$ | This study |
| RO235 | V. cholerae E7946 Δ*lacZ::kan*, ICEVchInd5 Δ*brxC*::*frt* ΔfloR pRO62 $Sm^R$, $Tm^R$, $Kan^R$, $Cm^R$ | This study |
| RO236 | V. cholerae E7946 Δ*lacZ::kan*, ICEVchInd5 Δ*brxC*::*frt* ΔfloR pRO77 $Sm^R$, $Tm^R$, $Kan^R$, $Cm^R$ | This study |
| RO243 | V. cholerae E7946 Δ*lacZ::kan*, ICEVchInd5 ΔfloR pRO62 SmR, TmR, KanR | This study |
| RO250 | V. cholerae E7946 ΔlacZ::Kan, ICEVchInd5 ΔfloR pRO11 $Sm^R$, $Tm^R$, $Kan^R$, $Cm^R$ | This study |
| RO251 | V. cholerae E7946 Δ*lacZ::kan*, ICEVchInd5 ΔfloR pKL06 $Sm^R$, $Tm^R$, $Kan^R$, $Cm^R$ | This study |
| RO335-336 | *V. cholerae* E7946 Δ*lacZ::kan*, ICE*Vch*Ban9 Δ*brxU*::*frt-spec* ΔfloR $Sm^R$, $Tm^R$, $Kan^R$, $Spec^R$ | This study |
| RO389 | *V. cholerae* E7946 Δ*lacZ::kan*, ICEVchInd5 Δ*brxC*::*frt* ΔfloR pRO37 $Sm^R$, $Tm^R$, $Kan^R$, $Cm^R$ | This study |
| RO390-391 | *V. cholerae* E7949 Δ*lacZ::kan*, ICE*Vch*Ban9 Δ*brxC*::*frt-spec* ΔfloR $Sm^R$, $Tm^R$, $Kan^R$, $Spec^R$ | This study |
| RO394 | *V. cholerae* E7949 Δ*lacZ::kan*, ICE*Vch*Ban9 Δ*brxC*::*frt* ΔfloR $Sm^R$, $Tm^R$, $Kan^R$ | This study |
| RO395 | *V. cholerae* E7949 Δ*lacZ::kan*, ICE*Vch*Ban9 Δ*brxC*::*frt* ΔfloR pRO11 $Sm^R$, $Tm^R$, $Kan^R$, $Cm^R$ | This study |
| RO396 | *V. cholerae* E7949 Δ*lacZ::kan*, ICE*Vch*Ban9 Δ*brxC*::*frt* ΔfloR pRO37 $Sm^R$, $Tm^R$, $Kan^R$, $Cm^R$ | This study |
| RO397 | *V. cholerae* E7949 Δ*lacZ::kan*, ICE*Vch*Ban9 Δ*brxC*::*frt* ΔfloR pKL06 $Sm^R$, $Tm^R$, $Kan^R$, $Cm^R$ | This study |
| RO554 | *V. cholerae* E7946 Δ*lacZ::kan*, ICE*Vch*Ban9 Δ*brxU*::*frt* ΔfloR $Sm^R$, $Tm^R$, $Kan^R$ | This study |
| Phage | Description | Reference |
| ICP1_2006_E | ICP1 phage isolate (accession number MH310934; referred to as ICP1) | [2] |
| ICP1_2006_E Δ*orbA* (ROΦ1) | ICP1_2006_E with a deletion in the *orbA* gene, referred to ICP1 Δ*orbA* | [1] |

[a] Streptomycin – *Sm*; Kanamycin – *Kan*; Spectinomycin – *Spec*; Chloramphenicol – *Cm*; Trimethoprim – *Tm*; Resistance – R; Sensitive – S

**Table S3: Plasmids**

| Plasmids | *E. coli* Strain | Genotype[a] | Reference |
|---|---|---|---|
| pCMB17 | CMB17 (*E. coli* S17) | Ptac-riboE-3xFLAG (codon optimized), *Cm^R^* | |
| pFlippase | SGH109 (*E. coli* S17) | Ptac-riboE-flippase, $Cm^R$ | [1] |
| pGp25(*orbA*) | KS2432 (*E. coli* S17) | Ptac-riboE-gp25 (*orbA*; (ADX87841.1)), $Cm^R$ | [1] |
| pKL06 | KS1870 (*E. coli* S17) | Ptac-riboE-empty vector, $Cm^R$ | [1] |
| pRO01 | RO87 (*E. coli* S17) | Ptac-riboE-*orbA-3xFLAG*, $Cm^R$ | This study |
| pRO02 | RO88 (*E. coli* S17) | Ptac-riboE-3xFLAG-*orbA*, $Cm^R$ | This study |
| pRO11 | RO106 (*E. coli* S17) | Ptac-riboE-*brxC*(*Vch*Ind5), $Cm^R$ | This study |
| pRO22 | RO151 (*E. coli* S17) | Ptac-riboE-*brxC-3xFLAG* (*Vch*Ind5), $Cm^R$ | This study |
| pRO24 | RO118 (*E. coli* S17) | Ptac-riboE-*brxC(K73A)* (*Vch*Ind5), $Cm^R$ | This study |
| pRO37 | RO383 (*E. coli* S17) | Ptac-riboE-*brxC* (*Vch*Ban9), $Cm^R$ | This study |
| pRO62 | RO231 (*E. coli* S17) | Ptac-riboE-$brxC^{E255A}$ (*Vch*Ind5), $Cm^R$ | This study |
| pRO76 | RO229 (*E. coli* S17) | Ptac-riboE-$brxC^{E255A}$-*3xFLAG* (*Vch*Ind5), $Cm^R$ | This study |
| pRO77 | RO233 (*E. coli* S17) | Ptac-riboE-$brxC^{K73A}$-*3xFLAG* (*Vch*Ind5), $Cm^R$ | This study |
| pRO97 | RO434 (*E. coli* XL1-Blue) | T-18-N-linker-*brxC* (*Vch*Ind5), $Amp^R$ | This study |
| pRO98 | RO435 (*E. coli* XL1-Blue) | T-18C-linker-*brxC* (*Vch*Ind5), $Amp^R$ | This study |
| pRO99 | RO436 (*E. coli* XL1-Blue) | T-25-N-linker-*brxC* (*Vch*Ind5), $Kan^R$ | This study |
| pRO100 | RO437 (*E. coli* XL1-Blue) | T-25-C-linker-*brxC* (*Vch*Ind5), $Kan^R$ | This study |
| pRO101 | RO414 (*E. coli* XL1-Blue) | T-18-N-linker-*orbA*, $Amp^R$ | This study |
| pRO102 | RO415 (*E. coli* XL1-Blue) | T-18-C-linker-*orbA*, $Amp^R$ | This study |
| pRO103 | RO416 (*E. coli* XL1-Blue) | T-25-N-linker-*orbA*, $Kan^R$ | This study |
| pRO104 | RO417 (*E. coli* XL1-Blue) | T-25-C-linker-*orbA*, $Kan^R$ | This study |
| pRO119 | RO455 (*E. coli* XL1-Blue) | T-18-N-linker-$brxC^{K73A}$(*Vch*Ind5), $Amp^R$ | This study |
| pRO120 | RO457 (*E. coli* XL1-Blue) | T-18-C-linker-$brxC^{K73A}$ (*Vch*Ind5), $Amp^R$ | This study |
| pRO121 | RO458 (*E. coli* XL1-Blue) | T-25-N-linker-$brxC^{K73A}$ (*Vch*Ind5), $Kan^R$ | This study |
| pRO122 | RO467 (*E. coli* XL1-Blue) | T-25-C-linker-$brxC^{K73A}$ (*Vch*Ind5), $Kan^R$ | This study |
| pRO123 | RO460 (*E. coli* XL1-Blue) | T-18-N-linker-$brxC^{E255A}$(*Vch*Ind5), $Amp^R$ | This study |

| | | | |
|---|---|---|---|
| pRO124 | RO462 (*E. coli* XL1-Blue) | T-18-C-linker-*brxC*$^{E255A}$ (*Vch*Ind5), *Amp*$^{R}$ | This study |
| pRO125 | RO463 (*E. coli* XL1-Blue) | T-25-N-linker-*brxC*$^{E255A}$ (*Vch*Ind5), *Kan*$^{R}$ | This study |
| pRO126 | RO465 (*E. coli* XL1-Blue) | T-25-C-linker-*brxC*$^{E255A}$ (*Vch*Ind5), *Kan*$^{R}$ | This study |
| pRO127 | RO507 (*E. coli* XL1-Blue) | T-18-N-linker-*brxC* (*Vch*Ban9), *Amp*$^{R}$ | This study |
| pRO128 | RO514 (*E. coli* XL1-Blue) | T-18-C-linker-*brxC* (*Vch*Ban9), *Amp*$^{R}$ | This study |
| pRO129 | RO508 (*E. coli* XL1-Blue) | T-25-N-linker-*brxC* (*Vch*Ban9), *Kan*$^{R}$ | This study |
| pRO130 | RO510 (*E. coli* XL1-Blue) | T-25-C-linker-*brxC* (*Vch*Ban9), Kan$^{R}$ | This study |
| pRO158 | RO518 (*E. coli* XL1-Blue) | T-18-N-linker-*brxC* (*Vch*Ind5) + *rbs-orbA*, *Amp*$^{R}$ | This study |
| pRO159 | RO520 (*E. coli* XL1-Blue) | T-18-C-linker-*brxC* (*Vch*Ind5) + *rbs-orbA*, *Amp*$^{R}$ | This study |

[a] Kanamycin – *Kan*; Chloramphenicol – *Cm*; Ampicillin – *Amp*; Resistance – R

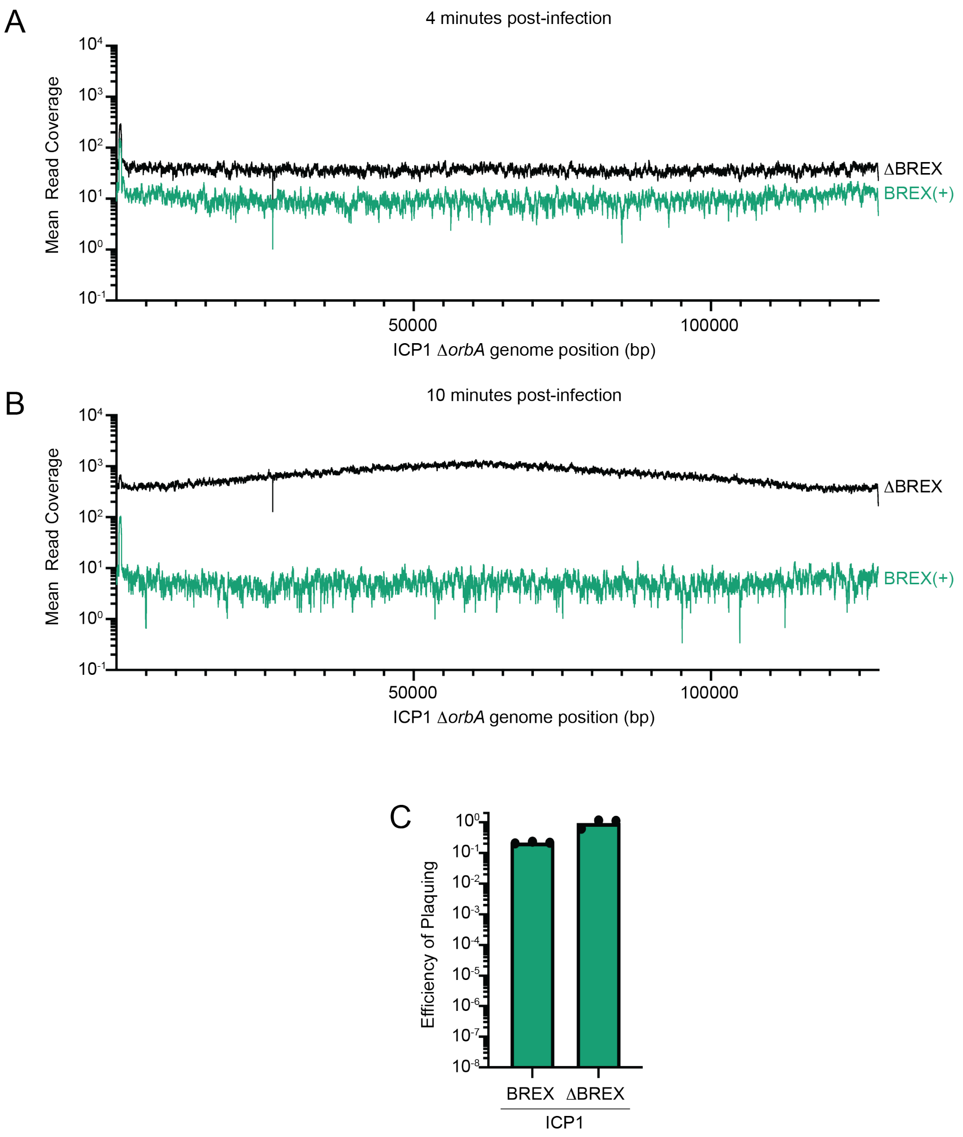


**Supplemental Figure 1. BREX restricts phage genome replication while OrbA restores phage plaquing. A-B)** Mean read coverage across the ICP1 Δ*orbA* genome in the presence and absence of BREX at **A)** 4 minutes post-infection and **B)** 10 minutes post-infection. *V. cholerae* cells either harboring the SXT *Vch*Ind5 (BREX(+); green) or where the BREX system and all of hotspot 5 were deleted (ΔBREX; black) were infected with ICP1 Δ*orbA*. The coverage shown is the mean of three biological replicates. **C)** Efficiency of plaquing (EOP) of ICP1 on *V. cholerae* that encodes the *Vch*Ind5 BREX system and deleted for the BREX system relative to the permissive background E7946 lacking *Vch*Ind5. Each dot represents a biological replicate.

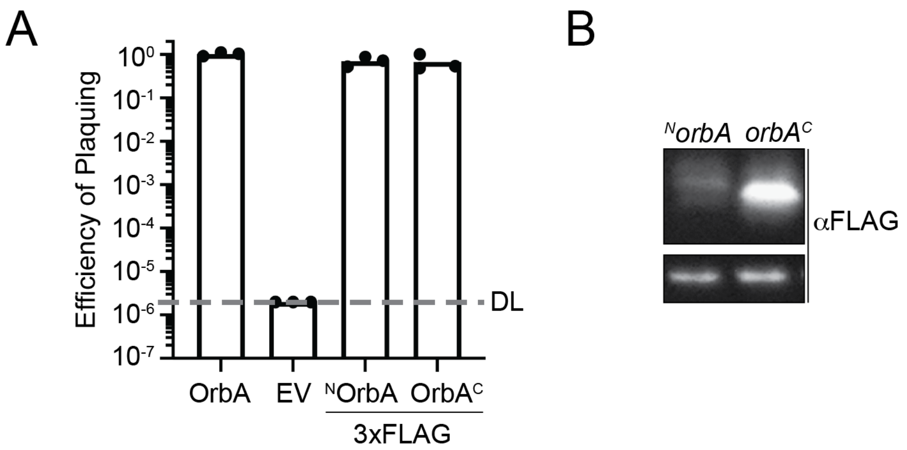


**Supplemental Figure 2. FLAG-tagged OrbA can inhibit BREX. A)** Efficiency of plaquing (EOP) of ICP1 Δ*orbA* on *V. cholerae* that encodes the *Vch*Ind5 BREX system carrying the vector indicated for expression of *orbA in trans* relative to the permissive strain *V. cholerae* E7946 lacking *Vch*Ind5 and harboring an empty vector. DL - detection limit. EV - empty vector. 3xFLAG translationally fused to the N-terminus of OrbA ($^N$OrbA). 3xFLAG translationally fused to the C-terminus of OrbA (OrbA$^C$) Each dot represents a biological replicate. **B)** Western blot analysis of FLAG-tagged OrbA using a primary antibody that recognizes the 3xFLAG tag. The bottom panel is a contaminating band from *V. cholerae* that is recognized by the FLAG antibody and is used as a loading control.

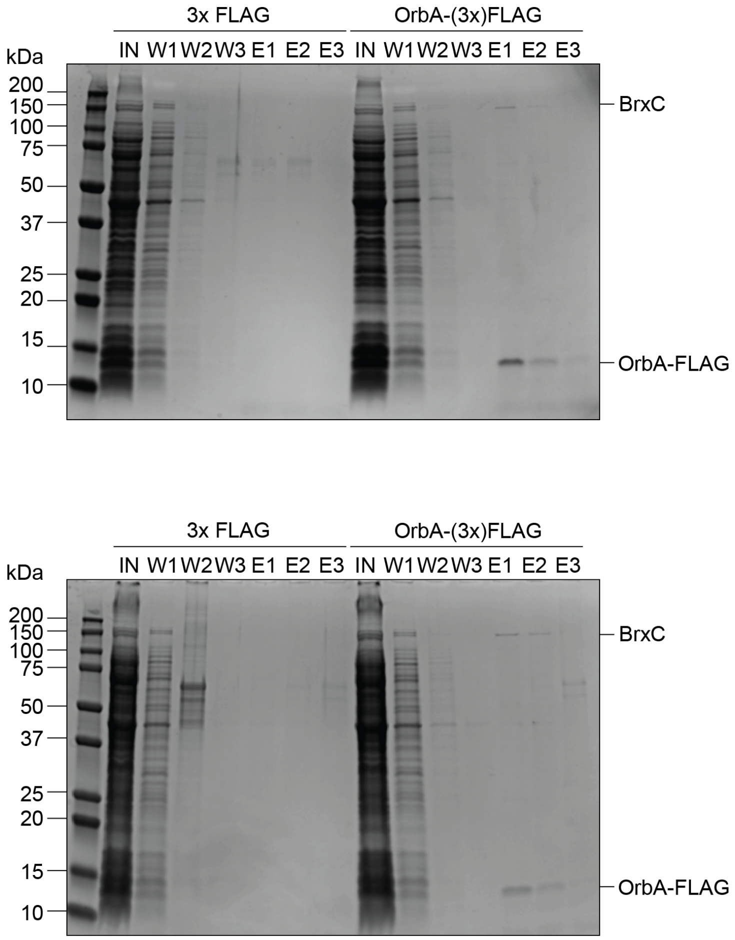


**Supplemental Figure 3.** Gel images of samples from replicates of 3xFLAG and OrbA-3xFLAG CoIPs resolved by SDS-PAGE and stained with Coomassie brilliant blue. IN – input. W – Wash. E – Elutions.

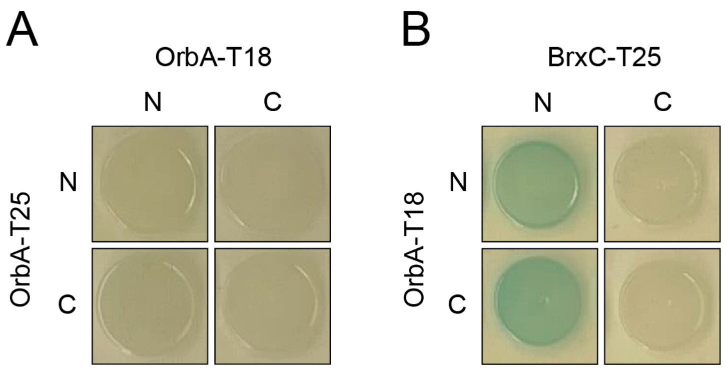


**Supplemental Figure 4.** OrbA interacts with BrxC. **A and B)** Bacterial two-hybrid analysis of the interaction between **A)** OrbA-OrbA and **B)** BrxC-OrbA. N- and C- indicate the terminus of each half of the adenylate cyclase enzyme was translationally fused to the protein of interest.

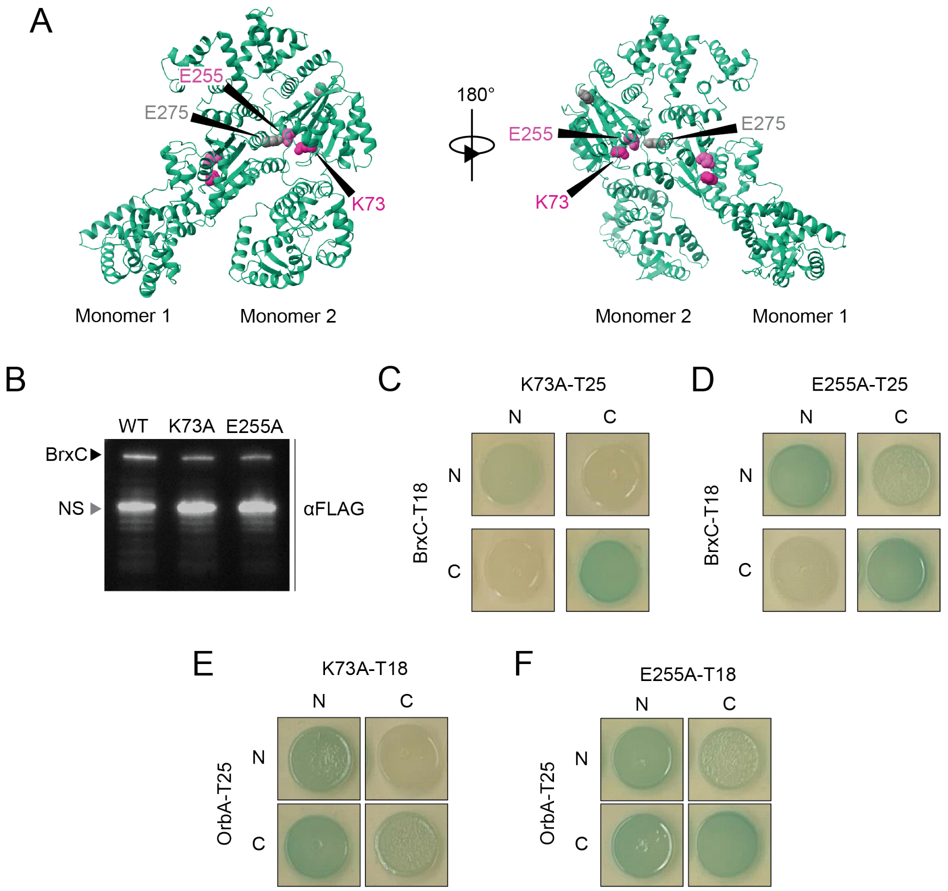


**Supplemental Figure 5. Mutation of the Walker A and B motif alters BrxC interactions. A)** Structural prediction of the N-terminal region of BrxC (1-532) as a dimer using ColabFold. Each BrxC monomer is indicated. The residues that were mutated in the Walker A and B motifs, K73 (dark pink) and E255 (light pink), respectively, are labeled on each monomer. The residue E275 (gray) was a candidate for mutation but did not have the features of a Walker B motif and was excluded from further study. **B)** Western blot analysis of either wild-type BrxC translationally fused to a C-terminal 3xFLAG tag or the Walker A (K73A) or Walker B (E255A) mutants using a primary antibody against the FLAG tag. NS - non-specific *V. cholerae* contaminating band used as a loading control. **C-F)** Bacterial two-hybrid analysis of the interaction between **C)** BrxC$^{WT}$-BrxC$^{K73A}$, **D)** BrxC$^{WT}$-BrxC$^{E255A}$, **E)** BrxC$^{K73A}$-OrbA, and **F)** BrxC$^{E255A}$-OrbA. N- and C- indicate the terminus of each half of the adenylate cyclase enzyme was fused to the protein of interest.

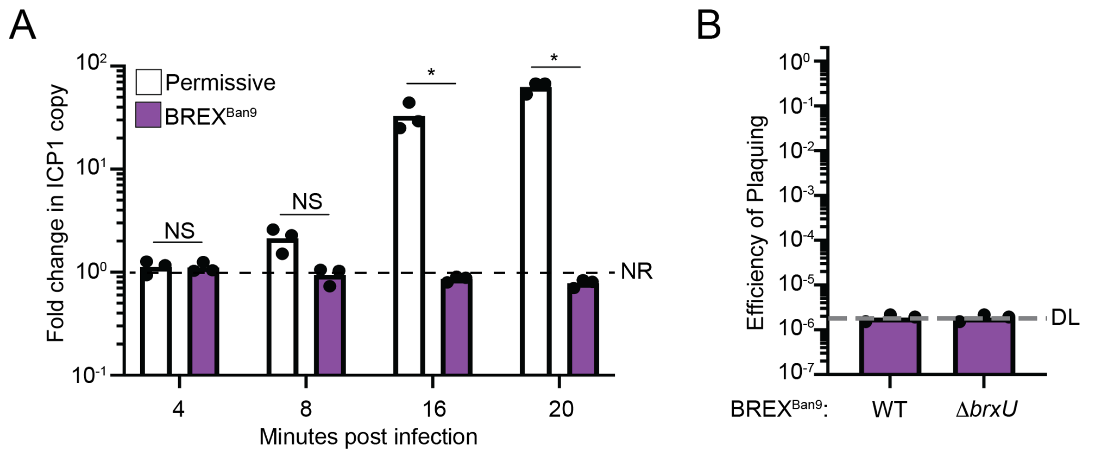


**Supplemental Figure 6. BrxU is not required for the inhibition of ICP1 by BREX$^{Ban9}$ A)** Quantification of ICP1 genome replication in the presence and absence of BREX$^{Ban9}$. Samples taken immediately following the addition of phage (Time 0) were compared to the indicated time points to calculate fold change. NR - no replication. Statistical analysis was performed using a student t-test was performed (* = $p<0.01$, ** = $p<0.001$, *** = $p<0.0001$, **** p=<0.00001). NS – not significant. Each dot represents a biological replicate **B)** Efficiency of plaquing (EOP) of ICP1 on *V. cholerae* either encoding a wild-type BREX system from *Vch*Ban9 (BREX$^{Ban9}$) or a BREX system from *Vch*Ban9 with *brxU* deleted (Δ*brxU*) relative to the permissive strain E7946 lacking *Vch*Ban9. DL - detection limit. Each dot represents a biological replicate.

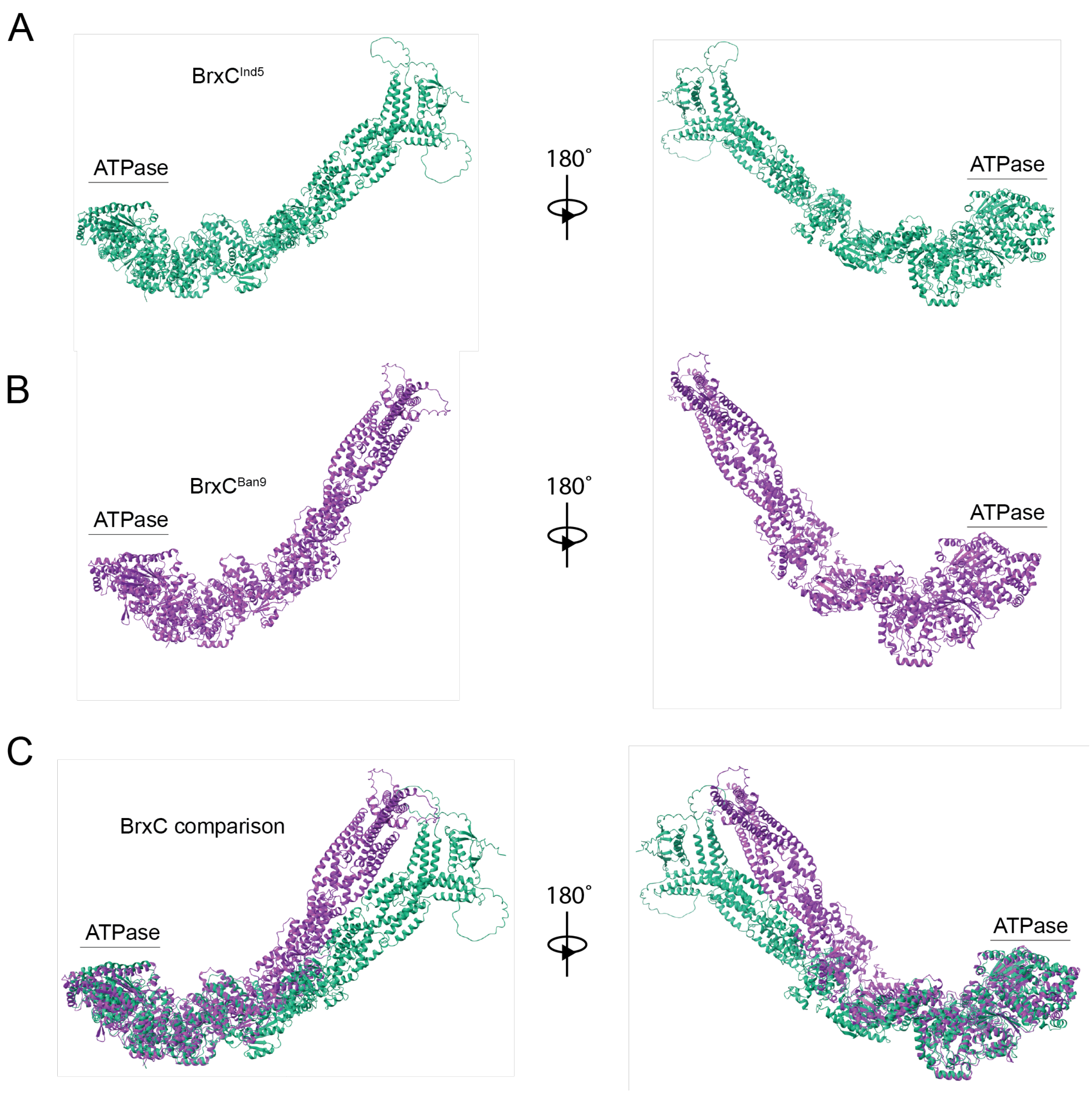


**Supplemental Figure 7. The predicted structure of BrxC$^{Ind5}$ is similar to BrxC$^{Ban9}$. A and B)** Predicted structures of BrxC encoded by **A)** *Vch*Ind5 (BrxC$^{Ind5}$) and **B)** *Vch*Ban9 (BrxC$^{Ban9}$) determined by ColabFold **C)** Structural comparison of the two BrxC proteins by Chimera. The N-terminal ATPase domain of both proteins is indicated.
